# Tracer-based lipidomics identifies novel disease-specific biomarkers in mitochondrial β-oxidation disorders

**DOI:** 10.1101/2023.08.31.555571

**Authors:** Marit Schwantje, Signe Mosegaard, Suzan JG Knottnerus, Jan Bert van Klinken, Ronald J Wanders, Henk van Lenthe, Jill Hermans, Lodewijk IJlst, Simone W Denis, Yorrick RJ Jaspers, Sabine A Fuchs, Riekelt H Houtkooper, Sacha Ferdinandusse, Frédéric M Vaz

**Author notes:** Shared first authors (Marit Schwantje, Signe Mosegaard). Shared last authors (Riekelt H Houtkooper, Sacha Ferdinandusse, Frédéric M Vaz).

## Abstract

Carnitine derivatives of disease-specific acyl-CoAs are the diagnostic hallmark for long-chain fatty acid oxidation disorders (lcFAOD), including carnitine shuttle deficiencies, very-long-chain acyl-CoA dehydrogenase deficiency (VLCADD), long-chain 3-hydroxyacyl-CoA dehydrogenase deficiency (LCHADD) and mitochondrial trifunctional protein deficiency (MPTD). The exact consequence of accumulating lcFAO-intermediates and possible influence on cellular lipid homeostasis are, however, still unknown. To investigate the fate and cellular effects of the accumulating lcFAO-intermediates and to explore new disease markers, we used tracer-based lipidomics with deuterium-labeled oleic acid (D9-C18:1) in lcFAOD patient-derived fibroblasts. In line with previous studies, we observed a trend towards neutral lipid accumulation in lcFAOD. In addition, we detected a direct connection between the chain length and patterns of (un)saturation of accumulating acylcarnitines and the various enzyme deficiencies. Our results also identified two new candidate disease markers. Lysophosphatidylcholine(14:1) (LPC(14:1)) was specifically increased in severe VLCADD compared to mild VLCADD and control samples. This was confirmed in plasma samples showing an inverse correlation with enzyme activity, which was better than the classic diagnostic marker C14:1-carnitine. The second biomarker is an unknown lipid class, which we identified as S-(3-hydroxyacyl)cysteamines. These are hypothesized to be degradation products of the CoA moiety of accumulating 3-hydroxyacyl-CoAs. S-(3-hydroxyacyl)cysteamines were significantly increased in LCHADD compared to controls and other lcFAOD, including MTPD. Our findings suggest extensive alternative lipid metabolism in lcFAOD and confirm that lcFAOD accumulate neutral lipid species. In addition, we present two new disease markers for VLCADD and LCHADD, that may have significant relevance for disease diagnosis, prognosis, and monitoring.

## Introduction

Fatty acids are, together with glucose and amino acids, the main substrates used to maintain metabolic homeostasis. Especially during fasting and exercise, organs such as the liver, heart and skeletal muscle rely highly on mitochondrial long-chain fatty acid β-oxidation (lcFAO) for energy production^1^. In addition to the generation of adenosine triphosphate (ATP) via the citric acid cycle and respiratory chain, fatty acids and their degradation products are important building blocks for the biosynthesis of different (macro)molecules^2,3^.

Deficiency of any of the enzymes of the mitochondrial lcFAO system leads to impaired degradation of fatty acids and energy shortage. This group of inherited metabolic diseases are called the long-chain fatty acid β-oxidation disorders (lcFAOD) and comprise seven different disorders; carnitine palmitoyltransferase 1 deficiency (CPT1D), carnitine-acylcarnitine translocase deficiency (CACTD), carnitine palmitoyltransferase 2 deficiency (CPT2D), very long-chain acyl-CoA dehydrogenase deficiency (VLCADD) and deficiencies of the mitochondrial trifunctional protein (MTP) including, generalized MTP deficiency (MTPD), long-chain 3-hydroxyacyl-CoA dehydrogenase deficiency (LCHADD) and long-chain acyl-CoA thiolase deficiency (LCKATD). LcFAOD can present with cardiomyopathy, rhabdomyolysis, hypoglycemia and for LCHADD/MTPD also with peripheral neuropathy and/or pigmentary retinopathy.

Biochemically, lcFAOD are characterized by the accumulation of disease-specific acyl-CoAs, reflecting the enzyme deficiency. Measurement of concomitantly accumulating acylcarnitines^1^ in plasma/blood spots is a well-established and cheap diagnostic tool that also allows newborn screening (NBS) for lcFAOD, as implemented in many countries worldwide. The clinical spectrum of patients with lcFAOD detected through NBS is highly variable, ranging from severe neonatal and fatal disease, to mild myopathic and even asymptomatic phenotypes^4^. It remains difficult to predict the clinical phenotype after diagnosis through NBS, complicating clinical decision making. In recent years, measurement of lcFAO flux in fibroblasts has been introduced as a reliable disease severity marker for lcFAOD^5^. Unfortunately, the lcFAO flux assay is only performed in a limited number of laboratories worldwide and requires a skin biopsy followed by fibroblast culturing, making this assay laborious and limited to only a small number of patients.

Although the acylcarnitine pattern is the diagnostic hallmark for lcFAOD, the exact role of the accumulation of lcFAO-intermediates in the pathophysiology of lcFAOD is largely unknown. It is hypothesized that certain accumulating lcFAO-intermediates may have toxic effects on tissues and organs, which together with metabolic energy shortage lead to clinical symptoms^6,7^. How the accumulation of lcFAO-intermediates influences cellular lipid homeostasis and development of symptoms is poorly understood. A number of studies have shown increased intracellular lipid accumulation in muscle fibres in muscle biopsies from patients with MTPD and VLCADD^8–11^. Other studies in lcFAOD patient-derived skin fibroblasts, cardiomyocytes and stem cell-derived LCHADD retina cells indicate a selective rerouting of long-chain fatty acids (LCFA) into triacylglycerols (TG)^12–14^. Alatibi *et al*.^13^, who reported the first lipidomic analyses in fibroblasts derived from patients with three different lcFAOD (CPT2D, LCHADD and VLCADD) proposed that the accumulation of specific acyl-CoAs leads to changes in the complex lipid profile of cells, which probably contributes to the clinical symptoms in lcFAOD patients.

In the present study, we used tracer-based lipidomics using deuterium-labeled oleic acid (D9-C18:1) to further explore the fate of accumulating long-chain fatty acids and lcFAO-intermediates in fibroblasts of four different lcFAOD; i.e. CPT2D, LCHADD, MTPD and VLCADD. We here report a direct correlation between the accumulation of specific lcFAO-intermediates and the various enzyme deficiencies, and a more general accumulation of neutral lipid species such as TG. Moreover, tracer-based lipidomics allowed for the identification of two new disease markers, LPC(14:1) for VLCADD and S-(3-hydroxyacyl)cysteamines for LCHADD, which may have significant relevance for disease diagnosis, prognosis, and monitoring.

## Materials and Methods

### Fibroblasts culture

Skin fibroblasts from six healthy controls, three CPT2D patients, five LCHADD patients, three MTPD patients, six severe VLCADD patients (lcFAO flux ≤10%) and six mild VLCADD patients (lcFAO flux >10%) were included (Table 1). For the patient and control fibroblasts used for the tracer-based lipidomics, informed consent was obtained for research studies. Cells were cultured in Ham’s F-10 medium (Gibco, Cat. No. 11550043) supplemented with 10% fetal calf serum (Invitrogen) 25 mmol/L HEPES, 100 U/mL penicillin, 100 μg/mL streptomycin, and 250 μg/mL amphotericin in a humidified atmosphere of 5% CO_2_ at 37°C. In parallel, all cell lines were cultured in the presence of a tracer, 50 μM D9-C18:1 (Avanti Polar Lipids, Cat. No 861809O) and 25 μmol/L L-carnitine (Sigma-Aldrich, Cat. No. C0158). After 96 hours of incubation, the fibroblasts were harvested by trypsinization and pellets were used for lipid isolation.

**Table 1.**
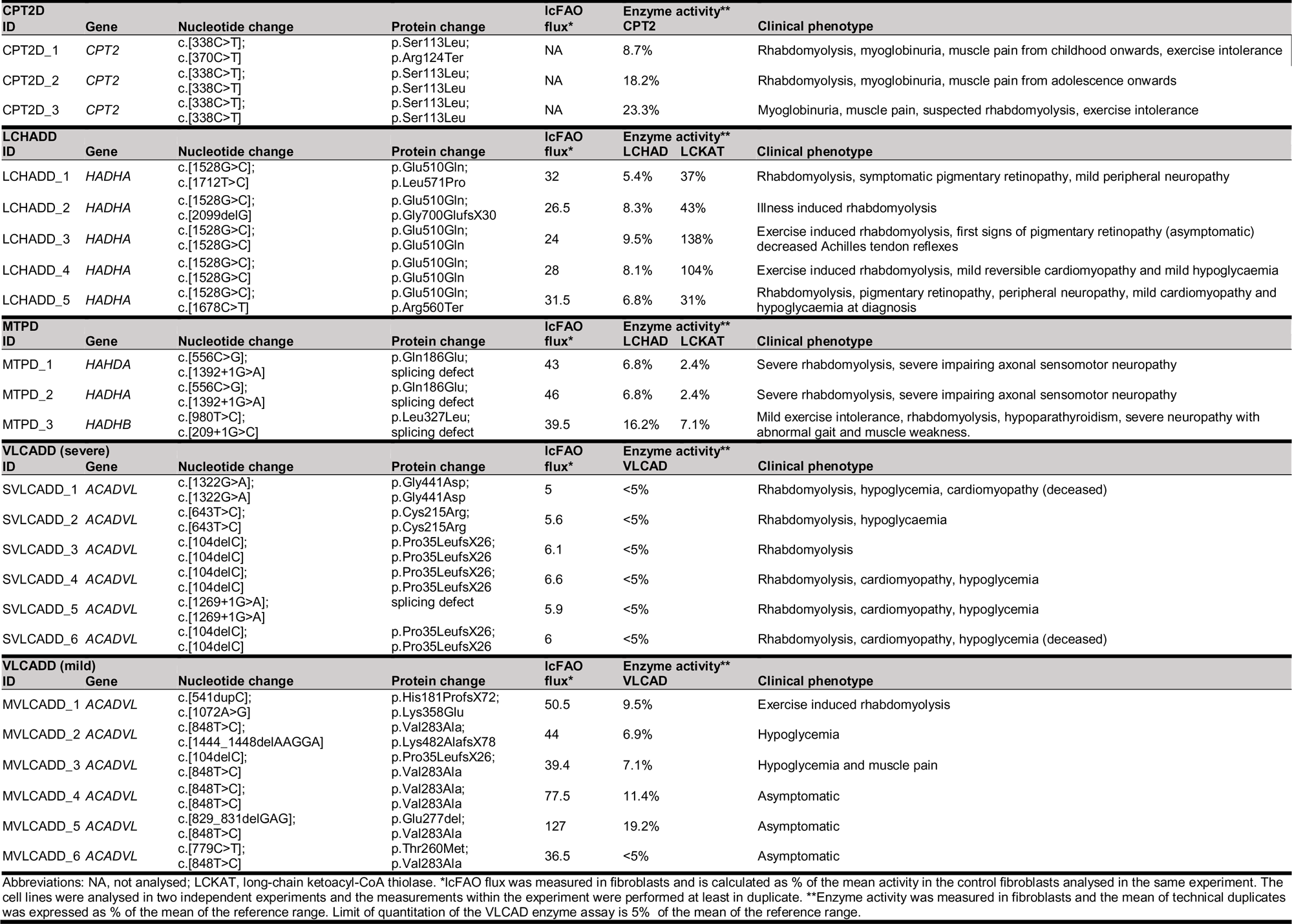
Patient cell lines characteristics.

### Single-phase lipidomic extraction

Fibroblast were resuspended in ultra-pure water and sonicated for 2 x 10 s at 8 W using a tip sonicator. For fibroblast used to exclude extracellular formation of detected lipid clusters, methanol:chloroform and internal standards were added before sonication. The protein concentration of the homogenates was determined with the BCA assay^15^ and 250 μg of protein equivalent was used for lipid extraction. The lipid extraction was performed in a 2 mL tube with 1.5 mL internal standards dissolved methanol:chloroform (1:1 (v/v)) as previously described with minor changes^16,17^. The following internal standards were added in listed quantities (per sample): Bis(monoacylglycero)phosphate BMP(14:0)_2_ (0.2 nmol), Ceramide-1-phosphate C1P (d18:1/12:0) (0.125 nmol), D7-Cholesteryl Ester CE(16:0) (2.5 nmol), Ceramide Cer(d18:1/12:0) (0.125 nmol), Ceramide Cer(d18:1/25:0) (0.125 nmol), Cardiolipin CL(14:0)_4_ (0.1 nmol), Diacylglycerol DAG(14:0)_2_ (0.5 nmol), Glucose Ceramide GlcCer(d18:1/12:0) (0.125 nmol), Lactose Ceramide LacCer(d18:1/12:0) (0.125 nmol), Lysophosphatidicacid LPA(14:0) (0.1 nmol), Lysophosphatidylcholine LPC(14:0) (0.5 nmol), Lysophosphatidylethanolamine LPE(14:0) (0.1 nmol), Lysophosphatidylglycerol LPG(14:0) (0.02 nmol), Phosphatidic acid PA(14:0)_2_ (0.5 nmol), Phosphatidylcholine PC(14:0)_2_ (2 nmol), Phosphatidylethanolamine PE(14:0)_2_ (0.5 nmol), Phosphatidylglycerol PG(14:0)_2_ (0.1 nmol), Phosphatidylinositol PI(8:0)_2_ (0.5 nmol), Phosphatidylserine PS(14:0)_2_ (5 nmol), Sphinganine 1-phosphate S1P(d17:0) (0.125 nmol), Sphinganine-1-phosphate S1P(d17:1) (0.125 nmol), Ceramide phosphocholines SM(d18:1/12:0) (2.125 nmol), Sphingosine SPH(d17:0) (0.125 nmol), Sphingosine SPH(d17:1) (0.125 nmol), Triacylglycerol TG(14:0)_3_ (0.5 nmol). The mixture was thoroughly mixed and sonicated in a water bath for 5 minutes followed by centrifugation at 4°C (16,000 × g for 5 minutes). Supernatant was transferred to a glass vial and evaporated under a stream of nitrogen at 60°C. The residue was dissolved in 150 μL of 1:1 (v/v) methanol:chloroform.

### Ultra performance liquid chromatography–high resolution mass spectrometry (UPLC– HRMS)

Lipids were analyzed using a Thermo Scientific Ultimate 3000 binary HPLC coupled to a Q Exactive Plus Orbitrap mass spectrometer. The normal phase separation of lipids was performed with 2 µL lipid extract that were injected on a Phenomenex, Luna 5µm Silica 100 Å, 250 mm x 2 mm maintained at 25°C. Mobile phase consisted of (A) 85:15 (v/v) methanol:water containing 0.025% formic acid and 6.7 mmol/L ammonia and (B) 97:3 (v/v) chloroform:methanol containing 0.025% formic acid. Using a flow rate of 0.3 mL/min, the LC gradient consisted of: Dwell at 10% A 0-1 min, ramp to 20% A at 4 min, ramp to 85% A at 12 min, ramp to 100% A at 12.1 min, dwell at 100% A 12.1-14 min, ramp to 10% A at 14.1 min, dwell at 10% A for 14.1-15 min.

For reverse phase separation of lipids, 5 μL lipid extract was injected onto an Acquity UPLC HSS T3 (100 x 2.1 mm, 1.8 μm, Waters) maintained at 60 °C. Mobile phase consisted of (A) 4:6 (v/v) methanol:water and B 1:9 (v/v) methanol:isopropanol, both containing 0.1% formic acid and 10 mmol/L ammonia. Using a flow rate of 0.4 mL/min, the LC gradient consisted of: Dwell at 100% A at 0 min, ramp to 80% A at 1 min, ramp to 0% A at 16 min, dwell at 0% A for 16-20 min, ramp to 100% A at 20.1 min, dwell at 100% A for 20.1-21 min. A Q Exactive Plus Orbitrap mass spectrometer (Thermo Fisher Scientific) was used in the negative and positive electrospray ionization mode using nitrogen as nebulizing gas. The spray voltage was 2500 V (neg) or 3500 V (pos), and the capillary temperature was 256°C. S-lens RF level: 50, Auxiliary gas: 11, Auxiliary gas temperature 300 °C, Sheath gas: 48 au, Sweep cone gas: 2 au. Mass spectra of lipids were acquired in both scan modes by continuous scanning from m/z 150 to m/z 2000 with a resolution of 280,000 full width at half maximum (FWHM) and processed using an in-house developed lipidomics pipeline^18^, written in the R programming language (http://www.r-project.org). The identified peaks were normalized to the intensity of the internal standard for each lipid class.

### LPC(14:1) measurements in lymphocytes

LPC(14:1) was analyzed using a Waters Acquity UPLC system coupled to a Waters Xevo TQ-S Micro mass spectrometer. The reversed phase separation was performed with 2 µL lipid extract that was injected on a Kinetex C8 LC Column (50 x 2.1 mm, 2.6 um, Phenomenex) maintained at 50°C. Mobile phase consisted of (A) water containing 0.1% formic acid and (B) methanol containing 0.1% formic acid. Using a flow rate of 0.4 mL/min, the LC gradient consisted of a linear gradient from 50% A at 0 min to 100% B at 8.35 min, 100% B from 8.35 to 12.5 min, to 50% A at 12.6 min and equilibrate at 50% A from 12.6-14.5 min. The TQ-S Micro mass spectrometer (Waters) was used in the positive electrospray ionization mode using nitrogen as nebulizing gas and argon as collision gas. The capillary voltage was 3500 V, the source temperature was 150°C and the desolvation temperature was 350°C. Cone gas flow 50 L/h an desolvation gas 650 L/hr. LPC(14:1) was measured in multiple reaction mode (MRM 466.3 > 104.1, cone 20 V, collision energy 23 V) and processed using Masslynx V4.2 (Waters). The peak area was normalized to the intensity of the internal standard D4-LPC(26:0) (MRM 640.5 > 104.1, cone 20 V, collision energy 30 V).

### Fragmentation analysis

Tandem mass spectrometry (MS/MS) analysis of the at first instance unknown lipid class was done using similar LCMS settings as described in the UPLC–HRMS section. MS/MS spectra were acquired in positive ion mode using higher energy collisional dissociation (HCD) using a normalized collision energy of 10, 35 and 60 eV for the m/z corresponding to the unknown lipid class. MS/MS spectra were evaluated using Thermo Scienctific Xcalibur 4.1.50.

### Annotation and statistical analysis

Peak identification in the untargeted lipidomics data was performed using a dedicated annotation tool written in MATLAB (https://www.mathworks/com), in which clusters of monoisotopic and labelled D9-lipid peaks could be annotated in a semi-automated fashion. Specifically, identification of known lipids was based on a combination of accurate mass and an in-house database of retention times from earlier studies and relevant standards. Identification of unknown lipid clusters was based on the visualization of all unannotated peaks with significant patient-control group differences and the subsequent comparison of within cluster mass differences with a pre-established list of commonly occurring mass differences that was composed of naturally occurring isotopes (e.g. ^13^C vs ^12^C), chemical moieties (e.g. CH_2_, H_2_, O) and adducts (e.g. [M+H]^+^ vs. [M+NH4]^+^).

Additional statistical analyses were performed using IBM SPSS Statistics for Windows, Version 26.0.0.1, Armonk, NY, IBM Corporation. Statistical significance of median differences between groups were calculated using two-tailed Mann-Whitney U-test. Correlation between to continuous variables was investigated using the two-tails Spearman correlation test. P values below 0.05 were considered statistically significant. Data are presented as median with range.

## Results

### Acylcarnitine patterns reflect enzyme deficiencies

Acylcarnitine levels in blood/plasma are generally thought to reflect intracellular levels of the corresponding acyl-CoA species. In line with this assumption, and with routine diagnostic procedures, we observed an increased relative abundance of disease-specific long-chain acylcarnitines (i.e., C18:1-carnitine in CPT2D, OH-C16:0-carnitine in LCHADD and MTPD, and C14:1-carnitine in VLCADD; Figure 1A) in lcFAOD patient fibroblasts cultured in standard HAM F-10 medium. When incubated with a medium containing deuterium-labeled oleic acid (D9-C18:1), we observed an increased relative abundance of D9-C18:1-carnitine, a derivative of the supplemented D9-C18:1 fatty acid, in all lcFAOD patients compared to control fibroblasts (Figure 1B) together with a decreased relative abundance of shorter chain intermediates in LCHADD and severe VLCADD patients (D9-C6:0- to D9-C10:0-carnitines; Figure 2A), indicating decreased β-oxidation capacity. Both the increased levels of disease-specific long-chain acylcarnitines and D9-C18:1-carnitine, and the decreased levels of short-chain D9-acylcarnitines are in line with the lcFAO enzyme deficiency in patient fibroblasts.

**Figure 1.**
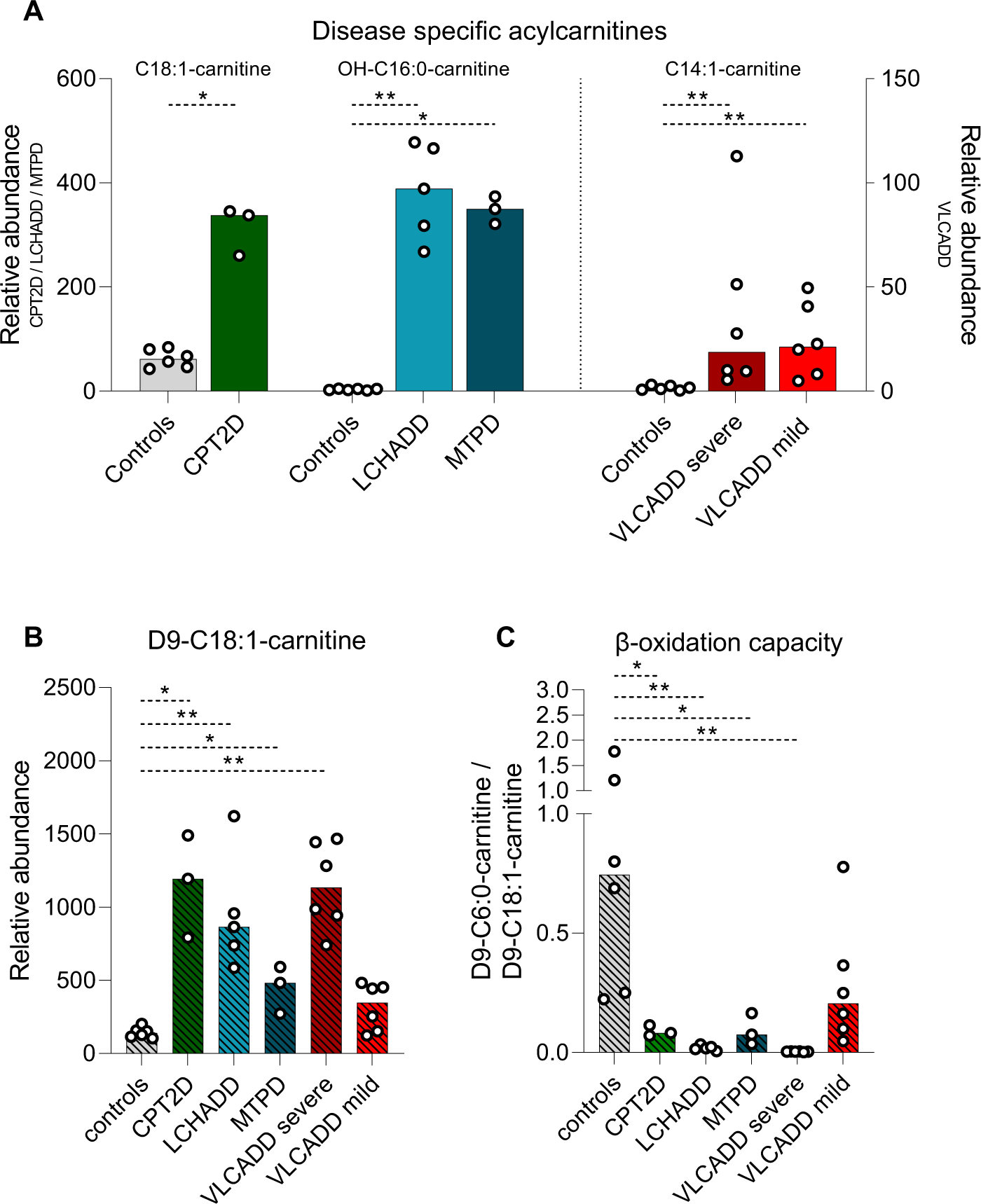
Disease-specific accumulation of acylcarnitines. A. The disease-specific accumulation of long-chain acylcarnitines in CPT2D (C18:1-carnitine), LCHADD and MTPD (OH-C16:0-carnitine), and VLCADD (C14:1-carnitine) fibroblasts and control fibroblasts under standard culture conditions. B. Increased relative abundance of D9-C18:1-carnitine 96 hours after addition of D9-C18:1 to the medium as a consequence of the impaired β-oxidation in all lcFAOD fibroblasts. C. The ratio of D9-C6:0-carnitine (represents product of β-oxidation) over D9-C18:1-carnitine (represents substrate of β-oxidation) as a measure of the remaining flux of D9-C18:1 through the β-oxidation system. *= p-value <0.05. **=p-value <0.01. Median (bars) and individual values (black circles) are shown in the graphs.

**Figure 2.**
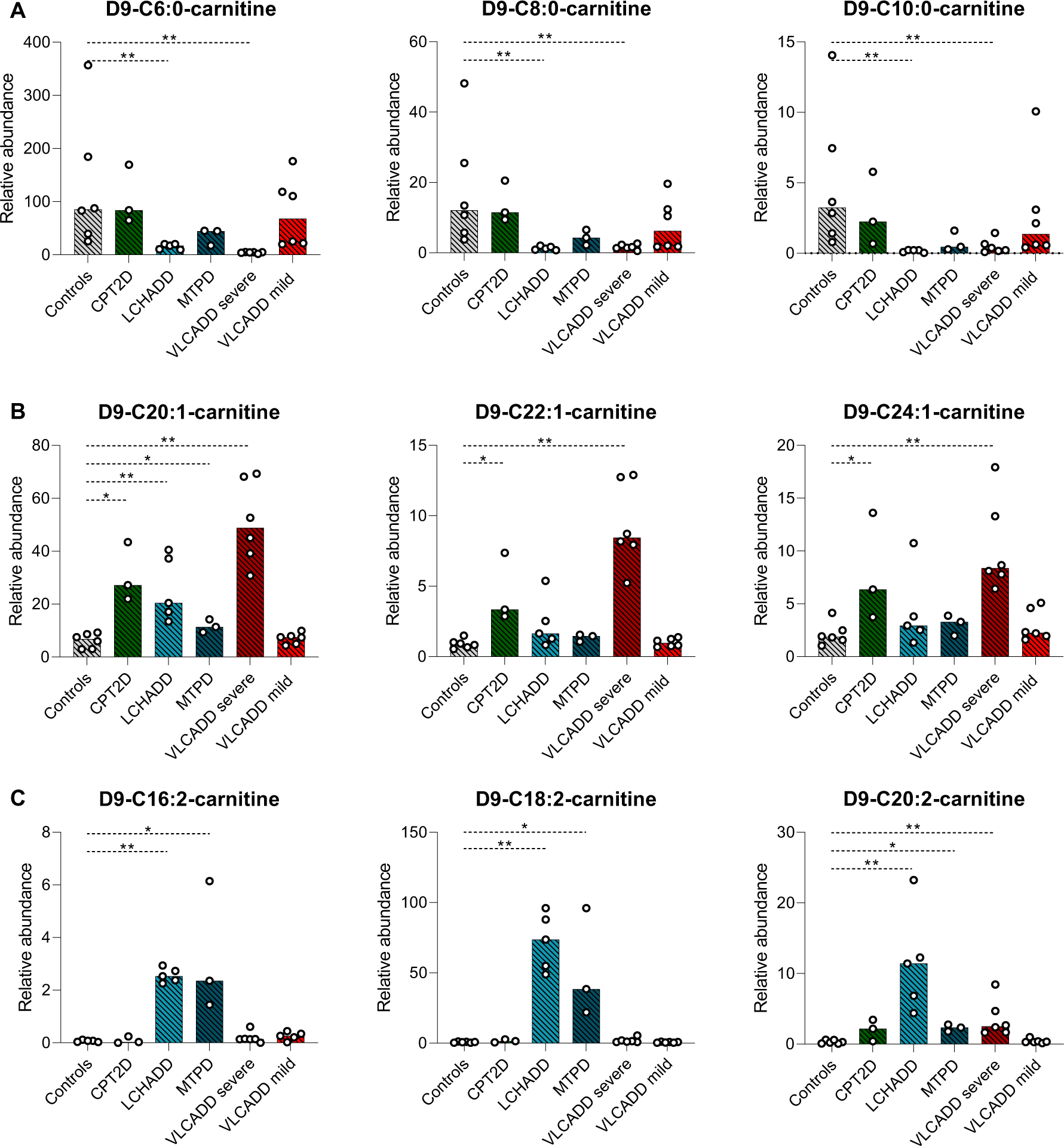
LcFAOD-specific handling of accumulating acylcarnitines. A. Relative abundance of shorter chain acylcarnitines (D9-C6:0- to C10:0-carnitine) in lcFAOD patient compared to control fibroblasts. B. Relative abundance of unsaturated and long-chain (>C18) D9-acylcarnitines in LCHADD and MTPD patient fibroblasts. C. Relative abundance of long-chain (>C18) acylcarnitines in lcFAOD patient fibroblasts compared to controls. *= p-value <0.05. **=p-value <0.01. Median (bars) and individual values (black circles) are shown in the graphs.

The mitochondrial lcFAO capacity can be represented as the ratio of D9-C6-carnitine over D9-C18:1-carnitine, the product and the substrate, respectively. As expected, controls had the highest lcFAO capacity. In LCHADD and severe VLCADD lcFAO capacity was severely impaired, but in mild VLCADD, CPT2D and MPTD there was residual lcFAO (Figure 1C). For VLCADD, this ratio correlated well with the lcFAO flux as measured with tritium-labeled oleic acid, a validated diagnostic tool to measure the flux of oleic acid through the β-oxidation system and a good predictor of the VLCADD phenotype^5^ (Spearman’s ρ = 0.85, p<0.001, Supplementary Figure 1A).

### Disease-specific lipid metabolism as identified by tracer-based lipidomics

When incubating CPT2D and VLCADD patient fibroblasts with D9-C18:1, we observed an increased relative abundance of (very) long-chain, mono-unsaturated acylcarnitines (D9-C20:1-carnitines and longer) compared to controls, indicating elongation of D9-C18:1 (Figure 2B). In LCHADD and MTPD, lipidomics analysis using normal phase liquid chromatography identified an increase of long-chain acylcarnitines with two double bonds, namely of D9-18:2-carnitine, D9-20:2-carnitine, and, with a lower relative abundance, D9-C16:2-carnitine (Figure 2C). We detected two separate peaks in the extracted-ion chromatograms of C18:2-carnitine and D9-C18:2-carnitine (Figure 3A and B). The first eluting peak (retention time (RT): 375-390s) was present in all lcFAOD, whereas the later eluting peak (RT: 390-430s) was only found in LCHADD and MTPD patient fibroblasts. This was also seen for C20:2-carnitine and C22:2-carnitine, although the relative abundances were increasingly lower (Figure 3C). We speculate that these LCHADD- and MTPD-specific lipid acylcarnitine species likely are carnitine esters that originate from the VLCAD-dependent dehydrogenation of D9-9-enoyl-C18-CoA (D9-)oleyl-carnitine and its elongation products D9-11-enoyl-C20-CoA and D9-13-enoyl-C22-CoA, resulting in the respective formation of D9-2,9-dienoyl-C18-CoA (D9-C18:2-CoA), D9-2,11-dienoyl-C20-CoA (=D9-C20:2-CoA) and D9-2,13-dienoyl-C22-CoA (=D9-C22:2-CoA). We observed a negative correlation between the calculated mitochondrial lcFAO capacity and the levels of D9-2,11-dienoyl-C20-carnitine accumulation in LCHADD and MTPD (Spearman’s ρ = −0.86, p=0.01, Supplementary Figure 1B). This negative correlation indicated that more D9-9-enoyl-C18-CoA is elongated and consequently more D9-2,11-dienoyl-C20-carnitine is formed in case of a lower mitochondrial lcFAO capacity caused by a more severe downstream metabolic block. These results were confirmed in patient fibroblasts cultured in standard medium, where we also observed increased levels of very long-chain acylcarnitines (>C22) in VLCADD compared to controls, and increased levels of double unsaturated acylcarnitines in LCHADD and MTPD compared to controls and other lcFAOD patient fibroblasts (Supplementary Figure 2).

**Figure 3.**
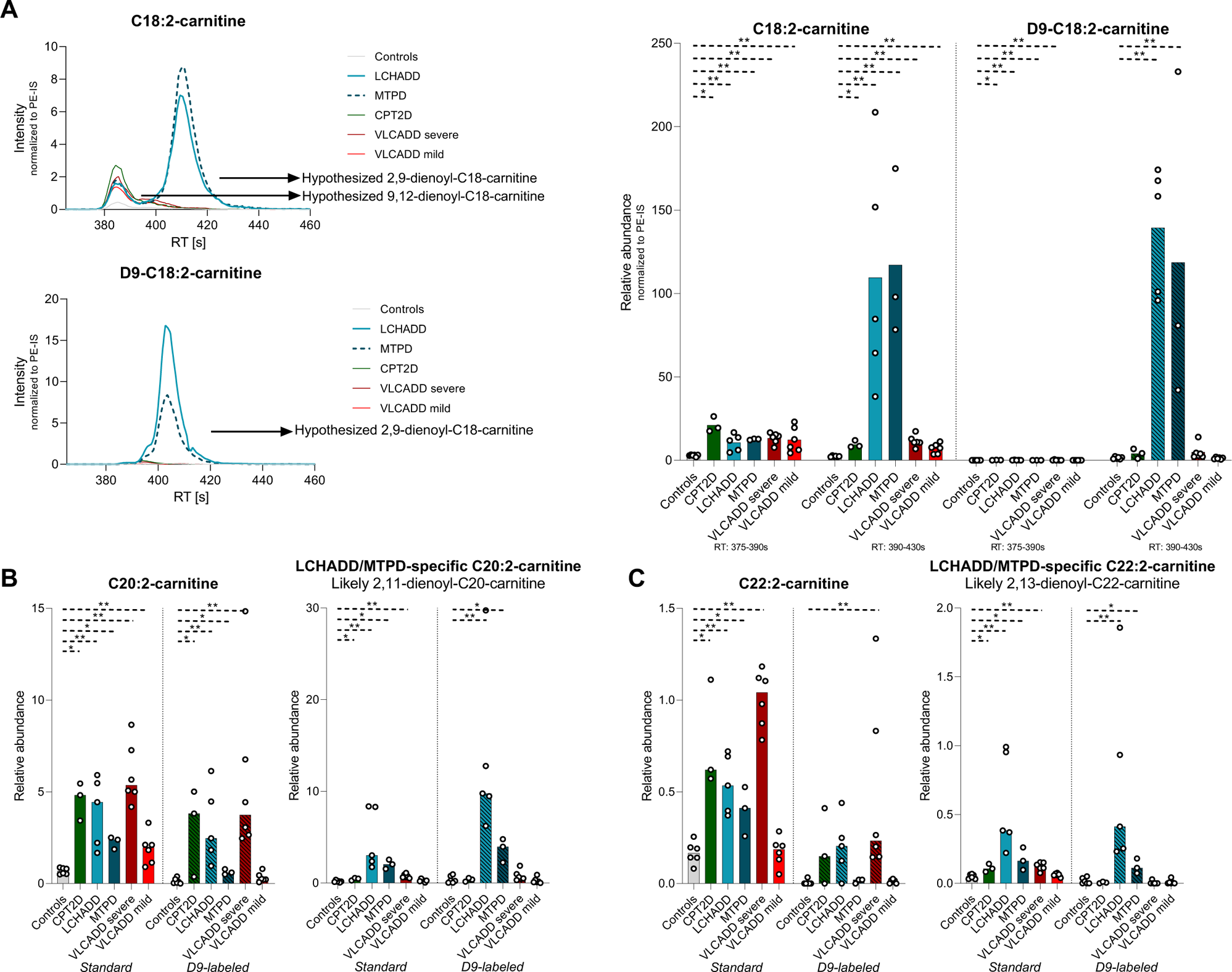
Extracted-ion chromatogram of LCHADD and MTPD-specific acylcarnitines. A. Extracted-ion chromatogram (left) and the peak areas of the signal intensity (normalized based on the PE internal standard) (right) of the two peaks of C18:2-carnitines measured with reverse phase liquid chromatography in control and lcFAOD patient fibroblasts cultured in standard medium and medium with D9-C18:1. For the different acylcarnitine species, the peaks of the D9-labeled acylcarnitines had a slightly lower RT, likely as a consequence of the nine deuterium atoms that are heavier than their non-labelled counterparts. B. Relative abundances of C20:2-carnitine (B) and C22:2-carnitine (C) and the LCHADD/MTPD-specific peaks, expected 2,11-dienoyl-C20 (A), and 2,13-dienoyl-C22-carnitines (C) measured with reverse phase liquid chromatography in controls and lcFAOD patient fibroblasts cultured in standard medium and medium with D9-C18:1. *= p-value <0.05. **=p-value <0.01. Median (bars) and individual values (black circles) are shown in the graphs.

### LCHADD- and MTPD-specific hydroxyacylcarnitine species

To investigate possible differences in the LCHADD- and MTPD-specific OH-C18:1- and OH-C18:2-carnitine species we evaluated their extracted-ion chromatograms derived from the reversed-phase analysis. For both OH-C18:1-carnitine and D9-OH-C18:1-carnitine, we detected one peak (RT: 366-398 sec), likely corresponding to the 3-hydroxyl form of (D9-)oleoyl-carnitine (Supplementary Figure 3). However, for OH-C18:2-carnitine, we detected three peaks (Figure 4). Like for (D9-)9-enoyl-C18-carnitine, (D9-)11-enoyl-C20-carnitine and (D9-)13-enoyl-C22-carnitine, the peaks of the labeled substrates were shifted to slightly lower retention times compared to the unlabeled substrates. We hypothesize that the first eluting peak (RT: 326-341 sec) corresponds to the 3-hydroxyl form of linoleoyl-carnitine (3-OH-9,12-dienoyl-C18-carnitine). The second and third eluting peak (RT 341-360 sec and 362-386 sec, respectively), were also found in the extracted-ion chromatogram of D9-labeled OH-C18:2-carnitine, indicating that both peaks were products of the added D9-C18:1. The second eluting peak was predominantly present in LCHADD- and to a lesser extent in MTPD-patient fibroblasts when compared to controls. We hypothesize that the second peak at RT 341-360 sec is a product of fatty acid desaturates (FADS)^19^, forming D9-5/6-enoyl-C18-CoA followed by partial oxidation through the lcFAO pathway to form D9-5/6-enoyl-3-OH-C18-CoA and its corresponding acylcarnitine, see discussion. The third, MTPD-specific, peak at RT 362-386 sec is most likely (D9-)3-oxo-C18:1-carnitine, the intermediate accumulating in case of LCKATD.

**Figure 4.**
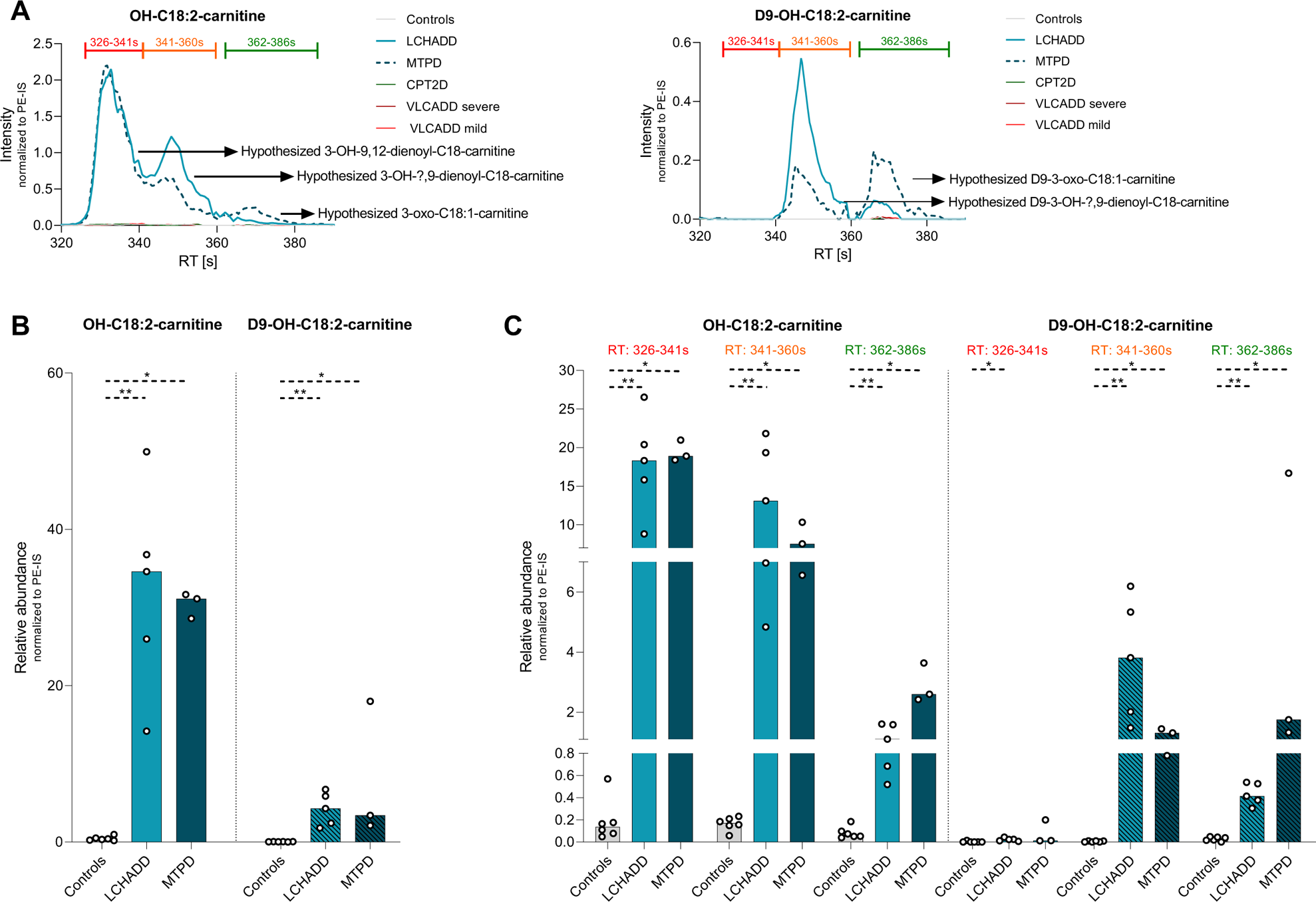
Different eluting peaks of OH-C18:2-carnitine. A. Extracted-ion chromatogram of OH-C18:2-carnitine (left) and D9-OH-C18:2-carnitine (right). For the OH-C18:2-carnitine, we observed three peaks. The peaks were hypothesized to be 3-OH-9,12-dienoyl-C18-carnitine (first peak, retention time (RT): 326-341 sec, red), 3-OH-?,9-dienoyl-C18-carnitine (second peak, RT: 341-360 sec, orange, position double bond unknown) and 2-oxo-C18:1-carnitine (third peak, RT: 362-386 sec, green). For the labeled analysis, D9-OH-C18:2-carnitine, we only observe two peaks, hypothesized to be D9-3-OH-?,9-dienoyl-C18-carnitine (first peak) and, D9-3-oxo-C18:1-carnitine (second peak). B. Relative abundance of OH-C18:2-carnitine and D9-OH-C18:2-carnitine in LCHADD and MTPD compared to controls. C. Relative abundance of OH-C18:2-carnitine and D9-OH-C18:2-carnitine for the different peaks (based on RT) observed in A. *= p-value <0.05. **=p-value <0.01. Median (bars) and individual values (black circles) are shown in the graphs.

### Neutral lipid accumulation in lcFAOD fibroblasts

In lcFAOD patient fibroblasts cultured in standard medium we observed a tendency towards neutral lipid accumulation. The sum of all triacylglycerols (TG), alkyldiacylglycerols (TG(O)) and cholesterol esters (CE) were 1.6, 1.9 and 2.9 times higher, respectively, in lcFAOD compared to control fibroblasts (Figure 5). Diacylglycerols (DG) and alkylacylglycerols (DG(O)) were only moderately increased, 1.2 and 1.3 times, respectively (Supplementary Figure 4). There were differences between the distinct lcFAOD: TG(O) and CE were mainly increased in LCHADD, and only mildly increased in other lcFAOD. CE levels were normal in severe VLCADD and MTPD (Figure 5C). Both the group of severe VLCADD patients with low lcFAO fluxes (5-6.6%), and the group of mild VLCADD patients with high lcFAO fluxes (33-125%) showed a tendency towards neutral lipid accumulation (Table 1, Figure 5). There was no statistically significant correlation between the amount of lipid accumulation (TG, TG(O) and CE) and lcFAO flux in VLCADD patients (not shown).

**Figure 5.**
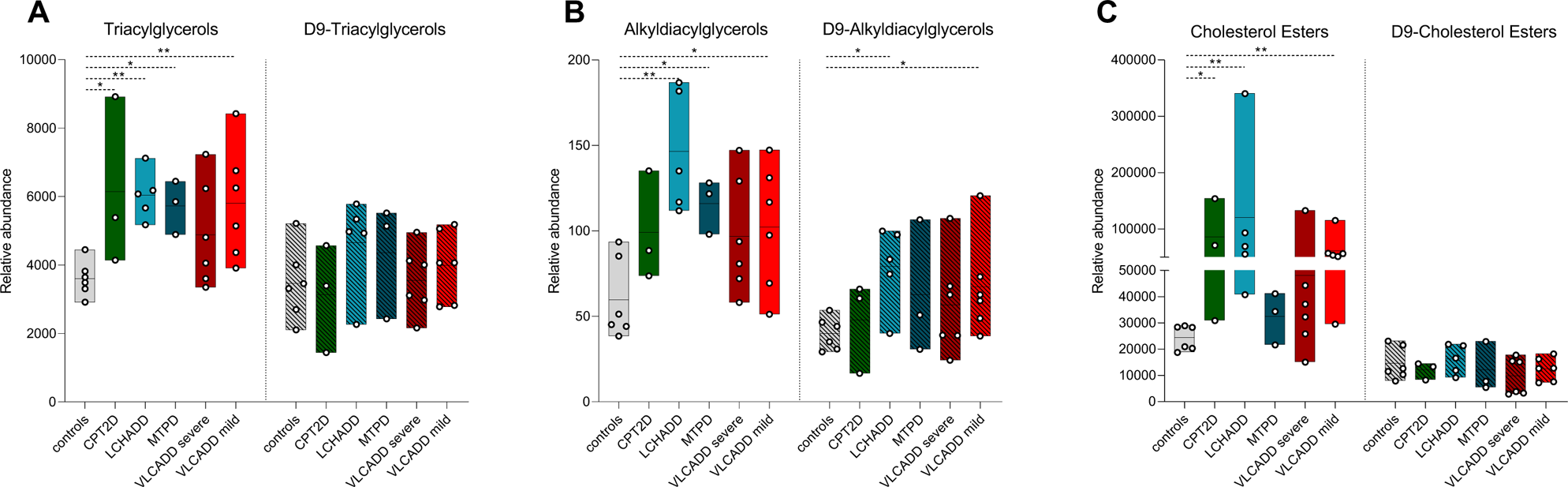
Neutral lipid species. The relative abundance of neutral lipids (TG, TG(O), and CE) in fibroblasts of most lcFAOD compared to controls under standard culture conditions and after adding D9-C18:1 to the medium. *= p-value <0.05. **=p-value <0.01. Median (black lines) and individual values (black circles) are shown in the graphs.

After D9-C18:1 loading to allow lipid tracing, the relative abundance of TG(O) was significantly increased in LCHADD and mild VLCADD, whereas D9-neutral lipids were not increased in other lcFAOD patient fibroblasts when compared to controls (Figure 5). Comparison of chain-length distribution and saturation patterns in neutral lipid chains with disease-specific acylcarnitine patterns did not show consistent effects (Supplementary Table 1). The relative abundance of phospholipids (phosphatidylcholine (PC), phosphatidylethanolamine (PE)) and lysophospholipids (lysophosphatidylcholine (LPC) and lysophosphatidylethanolamine (LPE)) was normal or slightly decreased compared to controls with and without D9-C18:1 added to the medium (Supplementary Figure 5). The ratio of PC and PE was comparable to the levels of the controls under both conditions (Supplementary Table 1). In conclusion, we observed a trend towards neutral lipid accumulation (TG, TG(O) and CE) in lcFAOD patient fibroblasts cultured in standard medium, whereas (lyso)phospholipids were not clearly affected.

### Identification of new disease markers for VLCADD

Although lysophopholipids as a group were not increased in lcFAOD patients, LPC(14:1) was significantly more abundant in severe VLCADD patient fibroblasts compared to controls (Figure 6A). This marker better discriminated between mild and severe VLCADD than the established VLCADD biomarker C14:1-carnitine and had a higher correlation with lcFAO-flux (Spearman’s ρ lcFAO flux and LPC(14:1):-0.82, p=0.001, Spearman’s ρ lcFAO flux and C14:1-carnitine: 0.13 p=0.70). Similarly, D9-LPC(14:1) had a significantly higher abundance in severe VLCADD compared to mild VLCADD and control fibroblasts (Figure 6A). To further investigate LPC(14:1) as a possible VLCADD disease severity marker, we measured LPC(14:1) in plasma of ten healthy controls, one carrier of VLCADD, and nine VLCADD patients with varying clinical and biochemical phenotypes (Supplementary Table 2). LPC(14:1) accumulation in plasma of VLCADD patients showed a strong correlation with enzyme activity measured in lymphocytes (r=-0.95, p<0.001)(Figure 6B). The correlation of LPC(14:1) with enzyme activity was better than the correlation of C14:1-carnitine with enzyme activity (Figure 6B Spearman’s ρ VLCAD-activity and C14:1-carnitine: −0.52, p=0.18, respectively). Correlating LPC(14:1) concentrations with clinical phenotype was complicated by the relatively low number of patients and the fact that this cohort contains both symptomatically- and NBS-diagnosed patients with varying follow-up times. Nevertheless, LPC(14:1) was increased in all symptomatic patients and one asymptomatic NBS-diagnosed patient with a low lcFAO flux (≤10%) associated with a severe clinical phenotype, whereas LPC(14:1) was normal compared to controls in one asymptomatic NBS-diagnosed patient with a high lcFAO flux.

**Figure 6.**
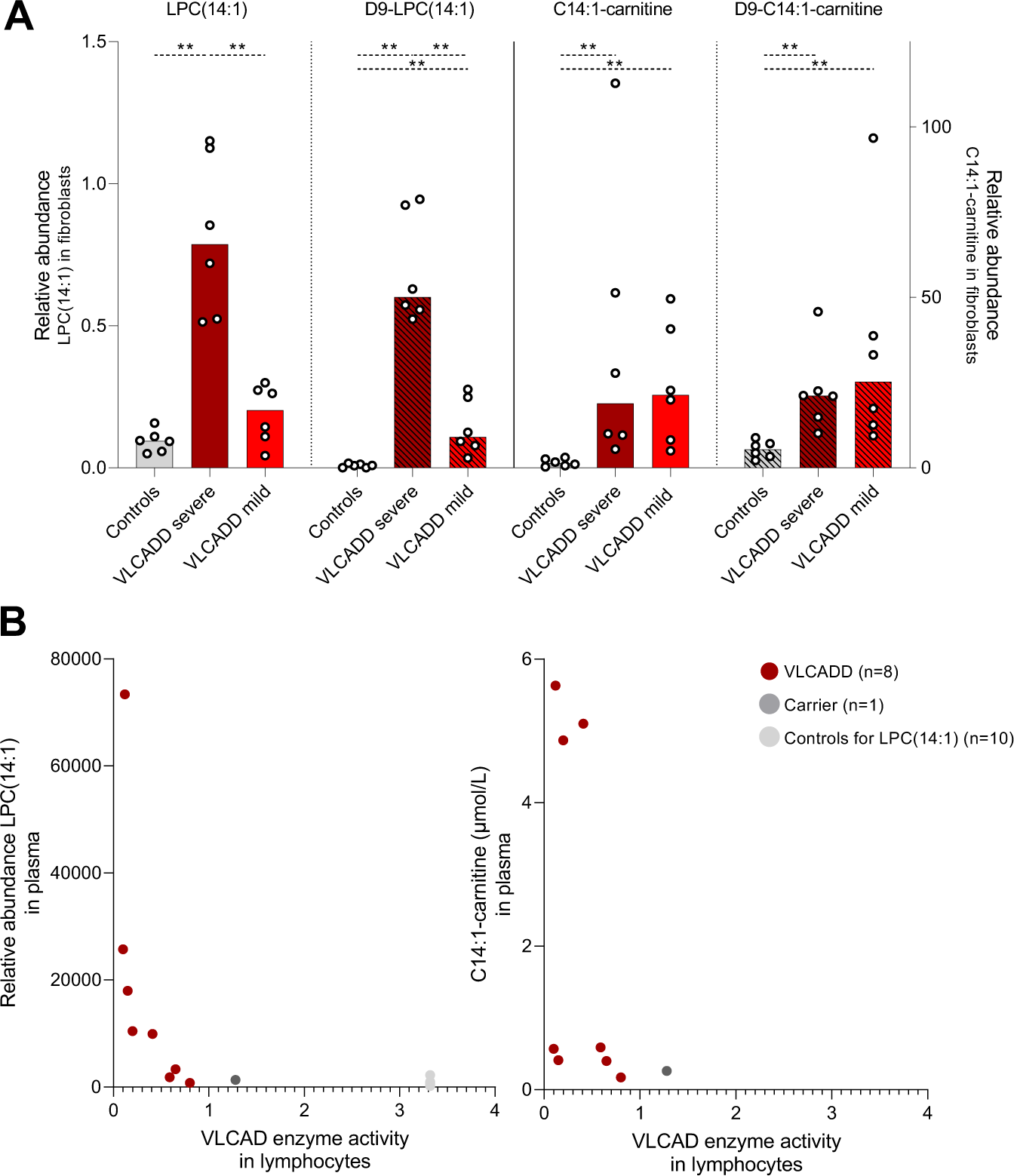
Disease severity marker for VLCADD. A. Relative abundance of LPC(14:1) and D9-LPC(14:1) in mild, severe VLCADD and controls. *= p-value <0.05. **=p-value <0.01. Median (bars) and individual values (black symbols) are shown in the graphs. B. LPC(14:1) and C14:1-carnitine were measured in plasma samples from healthy controls (n=10) (light grey circles), carrier of VLCADD (n=1) (dark grey circles), and VLCADD patients with varying phenotypes (n=8) (red circles).

### Identification of new disease markers for LCHADD

Using untargeted lipidomics, which allowed detection of ∼40.000 features, we identified a set of unknown features that were significantly elevated only in LCHADD when compared to controls and other lcFAOD patient fibroblasts (Figure 7A). The m/z values of the unknown features differed from each other by the mass of two methylene groups (CH_2_CH_2_, m/z difference: 28.0). After incubation with D9-C18:1, the D9-tracer was also incorporated in these, at that time unknown, features indicating that they are part of a lipid cluster containing fatty acids or their derivatives. Interestingly, in MTPD patient fibroblasts, this newly identified lipid cluster was not significantly increased. Fragmentation analysis was in line with the hypothesis that this unknown lipid cluster likely corresponded to thioesters of cysteamine, most likely originating from the degraded Coenzyme A-moiety, and a hydroxy-fatty acid (with varying chain lengths) (Supplementary Figure 6). We termed these molecules S-(3-hydroxyacyl)cysteamines.

**Figure 7.**
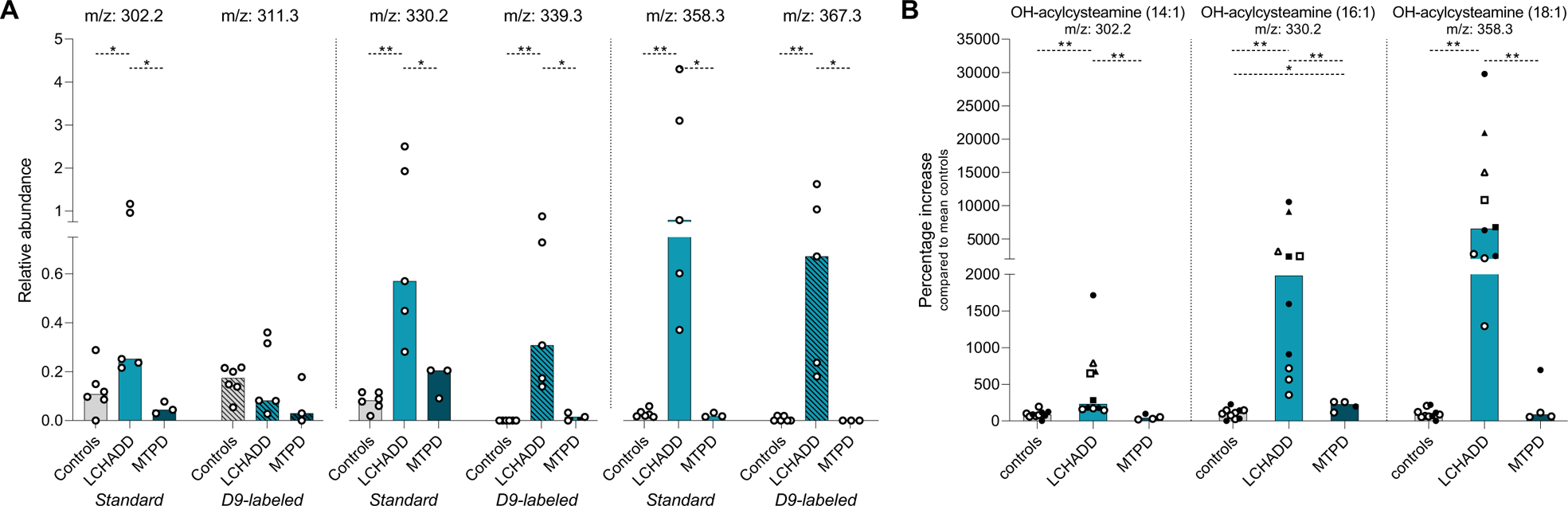
Disease specific marker for LCHADD. A. S-(3-hydroxyacyl)cysteamines (OH-acylcysteamine) were detected in significantly higher abundance in LCHADD patients compared to controls, and MTPD under standard culture conditions and after adding D9-C18:1 to the medium. *= p-value <0.05. **=p-value <0.01. B. Lipidomics analysis in fibroblast pellets from LCHADD patients (n=5) (two cell lines were included in the original lipidomics experiment: LCHADD_3 and LCHADD_4, Supplementary Table 3), an MTPD patient (n=1) and controls (n=4). Graphs show the combined results of the first (open circles) and the second (closed circles) lipidomics analysis. Median (bars) and individual values (circles) are shown as percentage of the controls’ average in the same experiment. LCHADD_3 and LCHADD_4 were also measured in the repeat experiment (analysis of new sample) and are shown in triangle and square symbols, respectively.

To confirm our findings, we repeated the lipidomics analysis in anonymized dermal fibroblast pellets not controlled for culture conditions and storage duration, from four healthy controls, one MTPD patient, and five LCHADD patients, again showing a significant increase in the LCHADD samples (Figure 7B). Next, to exclude that the S-(3-hydroxyacyl)cysteamines were an unspecific product produced extracellularly during sample preparation, fibroblasts of one control and one LCHADD patient were treated with methanol:chloroform before sonication to inactivate extracellular enzyme activities. S-(3-hydroxyacyl)cysteamines were similarly increased with and without methanol:chloroform treatment, with that excluding extracellular formation (Supplementary Figure 7).

## Discussion

Since the discovery of the first lcFAOD patients around the 1980s, diagnostic and therapeutic strategies have improved significantly, including implementation of lcFAOD in NBS programs allowing for pre-symptomatic diagnosis and treatment from birth. This has not only helped to prevent life-threatening hypoglycemia, but with time also showed that there still are difficulties to predict prognosis pre-symptomatically. Moreover, treatment strategies for some disorders are limited. Better understanding of the metabolic consequences of lcFAOD will likely contribute to further improving care for these patients, however, the fate and cellular effects of accumulating lcFAO-intermediates are poorly understood. In this study we used tracer-based lipidomics in CPT2D, LCHADD, MTPD and VLCADD patient fibroblasts to investigate the fate of long-chain fatty acids in lcFAOD, and the potential role of accumulating lcFAO intermediates in lcFAOD pathophysiology.

We observed a direct correlation between the chain length and saturation patterns of accumulating acylcarnitines with the enzyme deficiencies in lcFAOD. For CPT2D and VLCADD, we mainly detected accumulation of elongated mono-unsaturated acylcarnitines, whereas for LCHADD and MTPD, acylcarnitines with an additional double bond accumulated. Based on the combination of the low affinity of CPT2 for 2-enoyl-acylcarnitines^20^ and the accumulation of LCHADD/MTPD-specific long-chain acylcarnitines with an additional double bond we suggest that the corresponding activated fatty acids (e.g C18:1) were first elongated outside mitochondria (e.g. C20:1), then transported into mitochondria followed by dehydrogenation by VLCAD (e.g. C20:2).

In addition, tracer-based lipidomics and analysis of extracted-ion chromatograms of selected acylcarnitines, we found that there was alternative and disease-specific processing of accumulating lcFAO-intermediates. This is concluded from the finding of different chromatographic peaks in the extracted-ion chromatograms corresponding to different species of (D9-)OH-C18:2-carnitine in LCHADD and MTPD. Based on the labeling experiment we reasoned that the first and the third peak corresponded to the 3-hydroxyl form of linoleoyl-carnitine (3-OH-9,12-dienoyl-C18-carnitine) and 3-oxo-9-enoyl-C18 carnitine, respectively. The second peak, however, was only present in the labeling experiment and thus must have been derived from the D9-C18:1 which had gained an hydroxyl-group and an additional double bond to form (D9-)OH-C18:2 most likely introduced by the “general” desaturase FADS2^19,21^. We hypothesize that the (D9-)C18:1-CoA which now contains a second double bond introduced by the FADS enzyme (i.e. (D9-)9-enoyl-C18-CoA) to produce (D9-)C18:2-CoAs (position of double bond unknown), which is subsequently imported into mitochondria to be degraded via β-oxidation. Given the impairment in lcFAO because of LCHAD deficiency, β-oxidation is disturbed and (D9-)3-OH-C18:2-carnitine accumulates. These different species of OH-C18:2-carnitines illustrate the complex metabolic consequences of lcFAOD and highlight the insight that labeling studies in combination with UPLC-MS can yield.

Based on previous findings of neutral lipid accumulation due to impaired lcFAO both *in vitro* and *in vivo*^8–11,13,22^, we hypothesized that disease-specific lcFAO-intermediates are shunted towards neutral lipid species such as TG as a protection mechanism. Indeed, we observed a trend towards neutral lipid accumulation (TG, TG(O) and CE) in lcFAOD fibroblasts cultured in standard medium. (Lyso)phospholipids, however, did not accumulate, and in some lcFAOD the levels of these lipid species were even decreased. Moreover, we observed no changes in the PC/PE ratio, which was previously associated with mitochondrial disease^23–25^ and LCHADD^13^. The observed changes in neutral lipids but not in (lyso)phospholipids may relate to the different functions of these lipid species and likely is tissue/cell type-specific. Whereas neutral lipids mainly function as energy storage, (lyso)phospholipids are essential signaling molecules^26,27^ and building blocks for cellular and organellar membranes^28^. To prevent large changes in these structural/functional lipids, cells may shunt accumulating fatty acids into “storage” lipids as exemplified by steatosis of liver and muscle in many lcFAO^29–31^. However, tracing of D9-C18:1 did not provide direct evidence for storage of long-chain fatty acids in neutral lipids in lcFAOD fibroblasts. Supplementation of carnitine to the culture medium together with the D9-C18:1 may have stimulated detoxification through acylcarnitine formation and excretion in the medium. Also, the time-scale of the experiment could have been too short to accumulate D9-labeled fatty acids in neutral lipids.

C14:1-carnitine, the (NBS) marker for VLCADD, accumulated similarly in both severe and mild VLCADD patient fibroblasts. LcFAO flux measurements can discriminate between mild and severe VLCADD phenotypes^5^, but obtaining dermal fibroblasts for this is both invasive and time-consuming measurement with limited accessibility. We here found that LPC(14:1) also differentiated between severe and mild VLCADD in fibroblasts. Furthermore, we observed a correlation between LPC(14:1) in plasma from VLCADD patients and disease severity, but this initial observation does require confirmation in more patients. LPCs are the most abundant lysoglycerophospholipids in human blood, and are suggested to play, amongst others, a role in the inflammatory response with both pro- and anti-inflammatory effects^32–34^. LPCs can become toxic to cells at high concentrations, as they disrupt membrane structures and cause cell lysis, hence the prefix “lyso”. LPCs are derived from the corresponding phospholipid PC via hydrolysis by lipoprotein-associated phospholipases A_1_ or A_2_^34^. Long-chain LPCs are used as biomarkers for other metabolic diseases. For example, LPC(26:0) is used as a biomarker for adrenoleukodystrophy (ALD) and can be detected in bloodspots^35^. In VLCADD, accumulating C14:1 is likely integrated into PC eventually leading to the formation of LPC(14:1). As a product of PC, LPCs may be more stable and have a longer half-life in plasma than acylcarnitines, which are highly influenced by the patient’s diet, fasting and exercise^36^. If LPCs have a longer half-life, they might give a better representation of the accumulating intermediates over a longer time period which is less influenced by the exact sampling time. Although possible dietary and lifestyle influences on LPC(14:1) still need to be investigated, plasma LPC(14:1) may thus represent a useful marker to predict disease severity and monitor the effects of and compliance to dietary interventions.

Using untargeted lipidomics we identified a possible LCHADD-specific marker, which accumulated specifically in LCHADD patient fibroblasts in two separate experiments, including cell pellets of fibroblasts cultured under different conditions and at different time points. Based on the m/z values, fragmentation analysis, and the presence of a cluster of lipids with a fatty acid signature, we provisionally identified these lipids as S-(3-hydroxyacyl)cysteamines. We hypothesize that the generation of S-3-(hydroxyacyl)cysteamines occurs due to the high concentrations of 3-hydroxyacyl-CoAs that accumulate in LCHADD caused by the β-oxidation defect in combination with putatively lower affinity towards conversion into acylcarnitines (via CPT2) and cleavage into free fatty acids and coenzyme A (via thioesterases)^37^. The accumulating 3-hydroxyacyl-CoAs may be shunted into the CoA degradation pathway leading to the formation of S-(3-hydroxyacyl)cysteamines^38,39^. The exact mechanism of the generation of these lipid species is as yet unknown and requires further investigations. However, two enzymatic pathways/enzymes exist which in theory could produce S-(3-hydroxyacyl)cysteamines, pantheteinase^39^ and/or amidase^40,41^ activity. Pantheteinase (VNN) catalyzes the last step of the CoA degradation pathway but is generally regarded as an extracellular protein^39^. Accumulating 3-hydroxyacyl-CoAs could also be processed by an amidase, an enzyme that previously has been described to catalyze the once-step process in which acyl-CoAs are formed into acylcysteamines^40,41^.

We observed that S-(3-hydroxyacyl)cysteamines almost exclusively accumulated in LCHADD fibroblasts, whereas only minimal and inconsistent accumulation was observed in MTPD. Most of our MTPD patients had mild disease with relatively high residual enzyme activity and lcFAO capacity, likely allowing some degradation of 3-hydroxyacyl-CoAs via the β-oxidation pathway. Yet, it remains puzzling that hydroxyacylcarnitines accumulated in both LCHADD and MTPD, suggesting that hydroxyacyl-CoAs do so too, and that concomitantly S-(3-hydroxyacyl)cysteamines would be expected to be formed in both disorders.

Obviously, the LCHADD specificity of the S-(3-hydroxyacyl)cysteamines and the precise mechanism of formation still requires further investigation. However, our findings suggest that 3-hydroxyacyl-CoAs, and possibly also other accumulating acyl-CoAs, can be driven towards the CoA degradation pathway leading to the formation of S-((3-hydroxy)acyl)cysteamines. LCHADD and MTPD are unique among lcFAOD as they are the only two lcFAOD that display pigmentary retinopathy and peripheral neuropathy. For long, it was thought that toxic effects of accumulating 3-hydroxyacyl-CoAs and their carnitine derivatives may play a role in the development of these long-term complications. The exact pathophysiological mechanism however, remains unknown. Our discovery of the predominant and specific occurrence of S-(3-hydroxyacyl)cysteamines in LCHADD makes it tempting to speculate that they might play a role in LCHADD- (and MTPD-)specific disease pathophysiology. CoA biosynthesis deficiencies present with severe neurological phenotypes, including axonal neuropathy and pigmentary retinopathy due to CoA shortage^42,43,44^. We hypothesize that activity of the CoA degradation pathway in LCHADD and possibly MTPD may also result in CoA shortage which in turn might contribute to the development of pigmentary retinopathy and peripheral neuropathy.

In conclusion, we observed a tendency towards neutral lipid without (lyso)phospholipid accumulation in lcFAOD fibroblasts, suggesting protection of the intracellular balance in functional lipids by shunting accumulating fatty acids into neutral lipids. Tracer-based lipidomics did not only increase insight in the (alternative) processing and fate of accumulating lcFAO-intermediates in lcFAOD, but also revealed two new disease markers, LPC(14:1) for VLCADD and S-(3-hydroxyacyl)cysteamines for LCHADD, which may have significant relevance for disease prognosis and diagnosis, respectively.

## Data availability statement

Raw data included in the manuscript will be available upon reasonable requests.

## Author contributions

M.S., S.M., R.H., S.F., F.M.V., conceptualization; S.J.G.K., F.M.V., H.L., J.H., S.D., Y.R.J.J. methodology; J.B., M.S., S.M, formal analysis; M.S., S.M., S.J.G.K., H.L., J.H., S.D., Y.R.J.J. investigation; M.S., S.M., R.H., S.F., F.M.V., writing – original draft; M.S., S.M., R.J.W., S.A.F., R.H.H., S.F., F.M.V., writing – review & editing, M.S., S.M., F.M.V, visualization.

## Abbreviations

CACTD: carnitine-acylcarnitine translocase deficiency
CE: cholesterol esthers
CPT1D: carnitine palmitoyltransferase 1 deficiency
CPT2D: carnitine palmitoyltransferase 2 deficiency
DG: diacylglycerols
FADS: fatty acid desaturates
lcFAOD: long-chain fatty acid oxidation disorders
LCHADD: long-chain 3-hydroxyacyl-CoA dehydrogenase deficiency
LCKATD: long-chain acyl-CoA thiolase deficiency
LPC: lysophosphatidylcholine
LPE: lysophosphatidylethanolamine
MPTD: mitochondrial trifunctional protein deficiency
NBS: newborn screening
PC: phosphatidylcholine
PE: phosphatidylethanolamine
RT: retention time
TG: triacylglycerols
TG(O): alkyldiacylglycerols
VLCADD: very-long-chain acyl-CoA dehydrogenase

