## Supplementary Figures for "Tracer-based lipidomics identifies novel disease-specific biomarkers in mitochondrial β-oxidation disorders"

**Supplementary Figure 1**

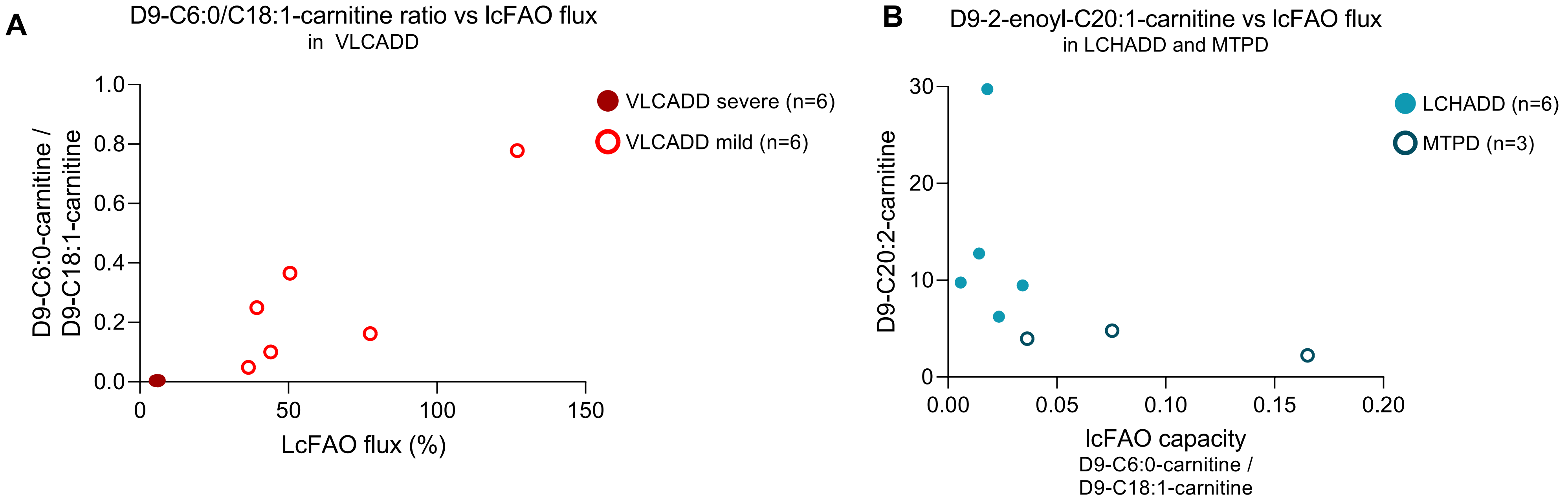

**Supplementary Figure 2**

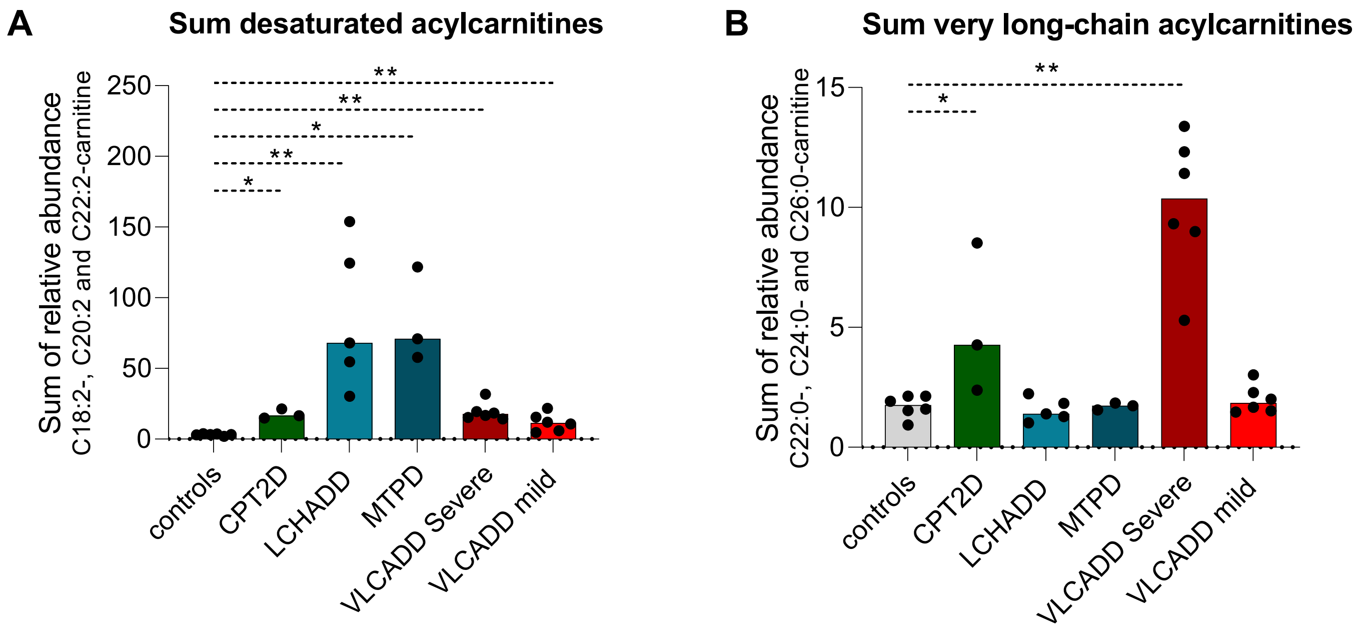

**Supplementary Figure 3**

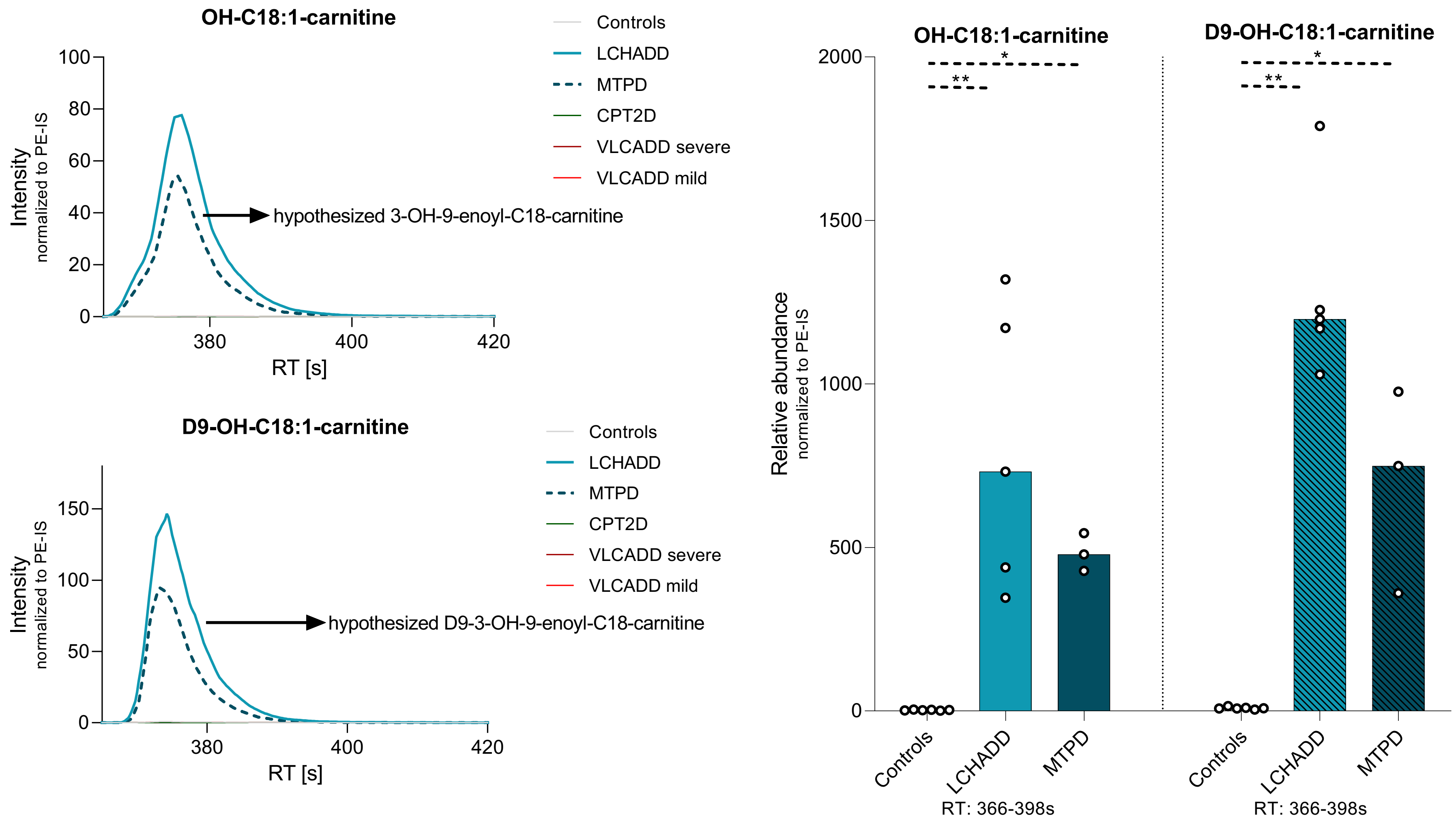

### Supplementary Figure 4

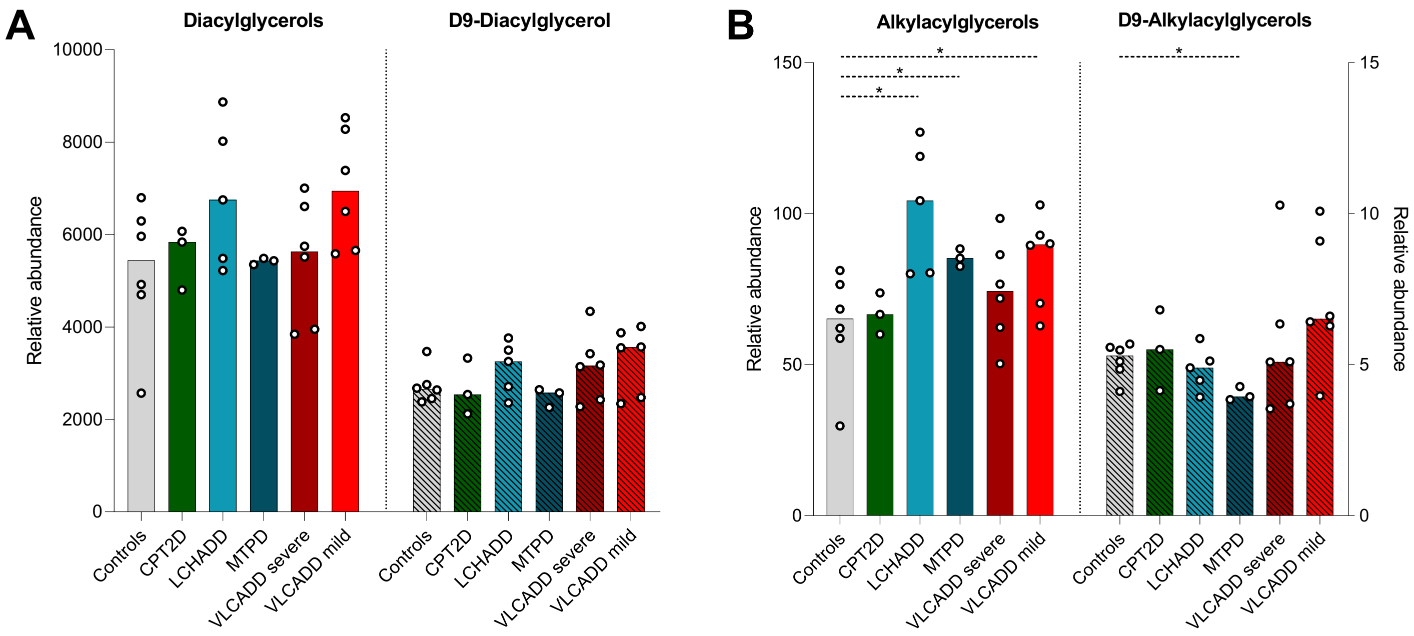

### Supplementary Figure 5

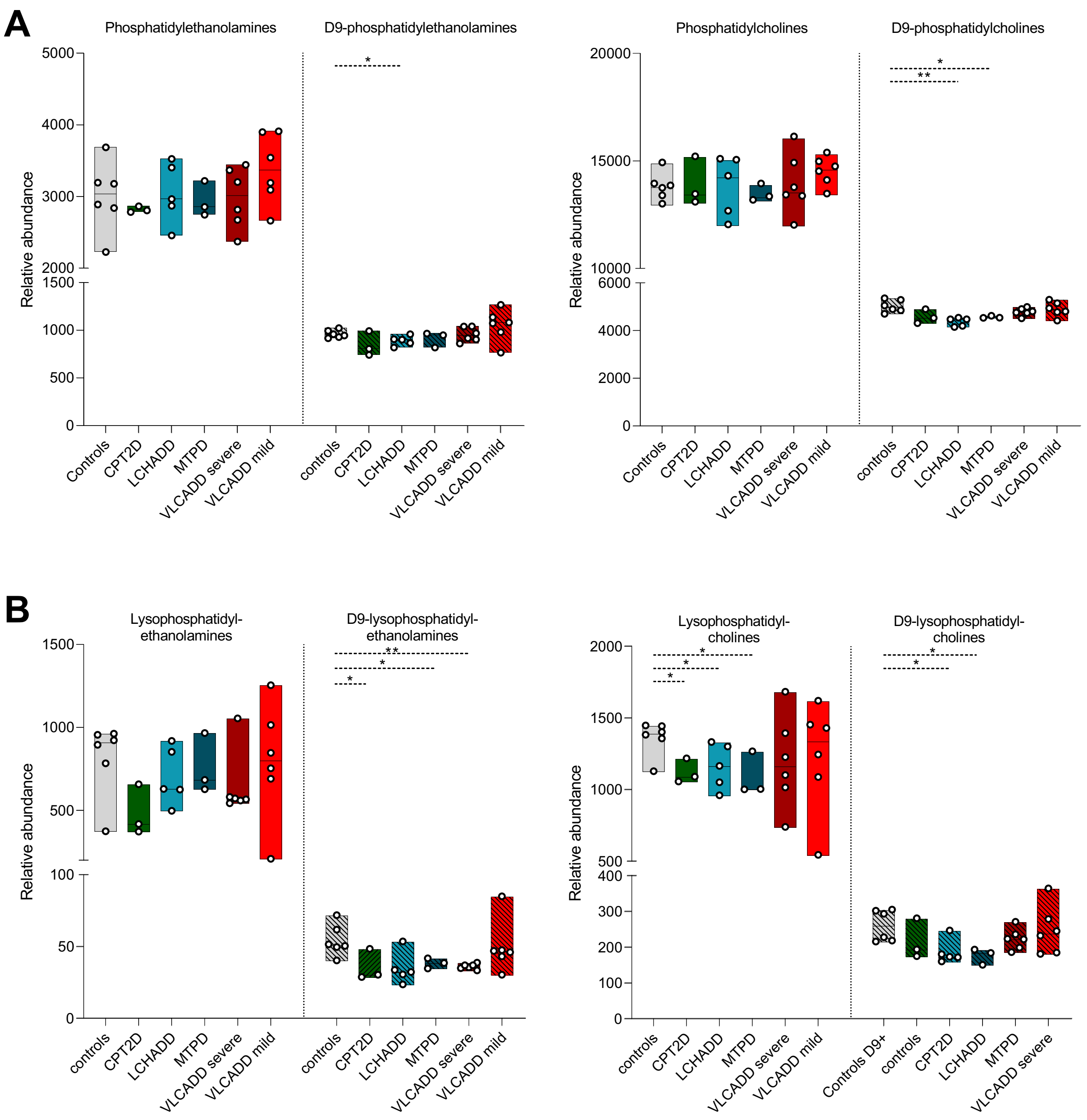

### Supplementary Figure 6

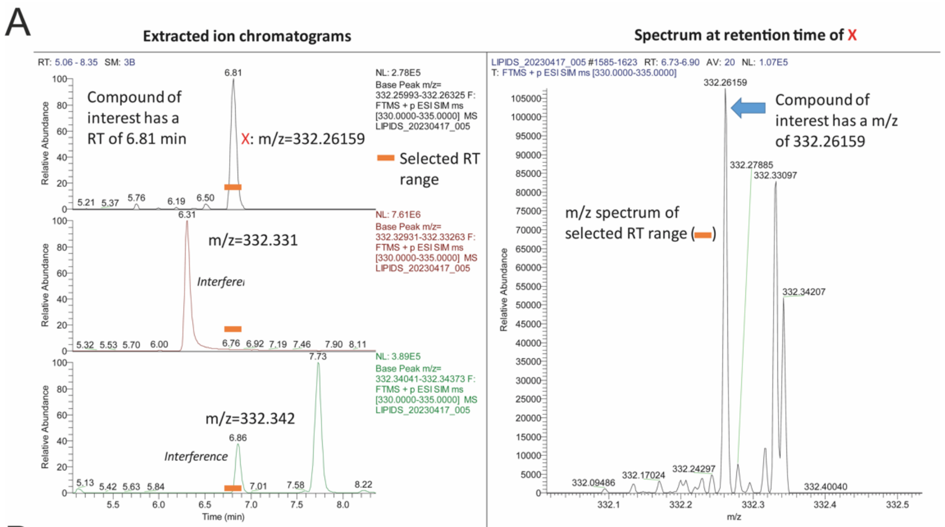


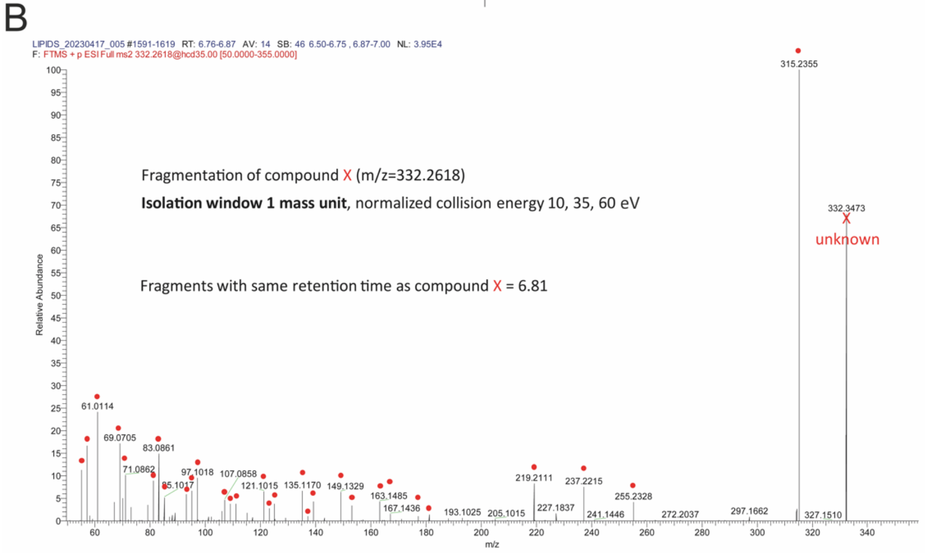


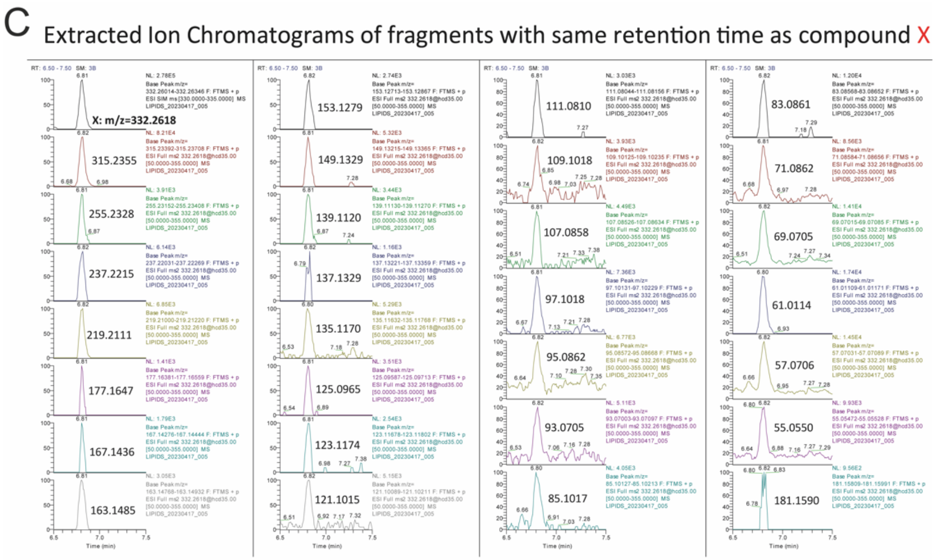


D Identification of fragments based on mass (delta mass < 0.001 mass unit) with correct retention time of 6.81

Red are fragments shown in structure below

| formula | m/z |
|---|---|
| C18H38O2NS+ | 332.2618 |
| C18H35O2S+ | 315.2355 |
| C16H31O2+ | 255.2328 |
| C16H29O+ | 237.2215 |
| C16H27+ | 219.2111 |
| C12H21O+ | 181.1590 |
| C13H21+ | 177.1647 |
| C11H19O+ | 167.1436 |
| C12H19+ | 163.1485 |
| C10H17O+ | 153.1279 |
| C11H17+ | 149.1329 |
| C9H15O+ | 139.1120 |
| C10H15+ | 135.1170 |
| C8H13O+ | 125.0965 |
| C9H15+ | 123.1174 |
| C9H13+ | 121.1015 |
| C7H11O+ | 111.0808 |
| C8H13+ | 109.1018 |
| C8H11+ | 107.0858 |
| C7H13+ | 97.1018 |
| C7H11+ | 95.0862 |
| C7H9+ | 93.0705 |
| C6H13+ | 85.1017 |
| C6H11+ | 83.0861 |
| C5H11+ | 71.0862 |
| C5H9+ | 69.0705 |
| C2H5S+ | 61.0114 |
| C4H9+ | 57.0706 |
| C4H7+ | 55.0550 |

Compound: X(16:0), m/z = 332.2618

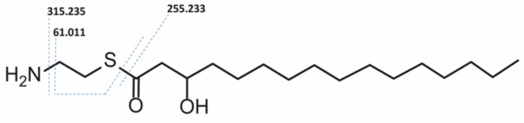


Other fragments likely result from further fragmentation of the aliphatic side chain

### Supplementary Figure 7

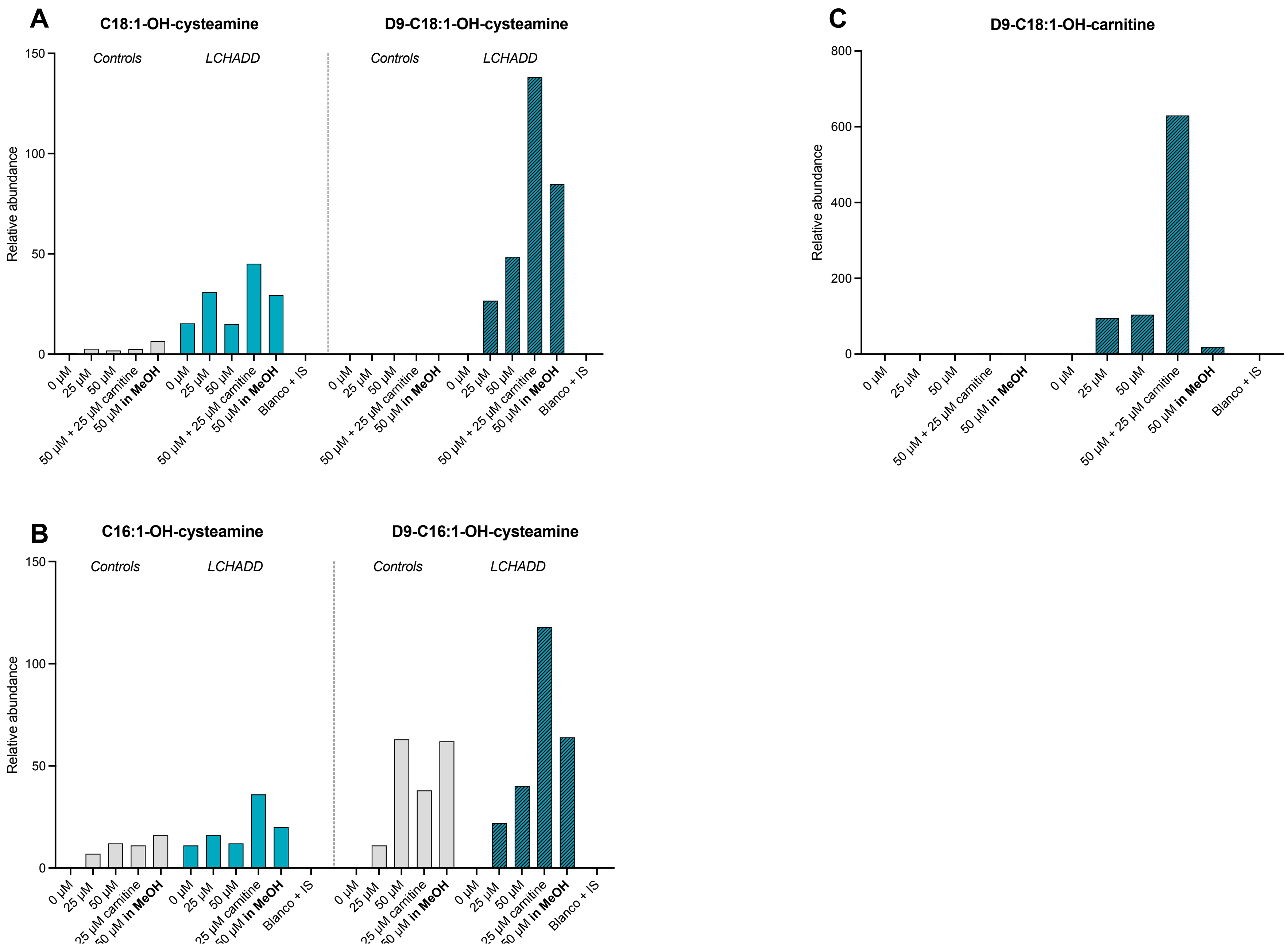
