## Supplementary tables and legends for "Tracer-based lipidomics identifies novel disease-specific biomarkers in mitochondrial β-oxidation disorders"

**Supplementary Table 1.**

**A. Distribution of saturation/chain length of neutral lipids**

| Lipids | Saturation (% of sum) | Controls n=6 median (range) | CPT2D n=3 median (range) | LCHADD n=5 median (range) | MPTD n=3 median (range) | VLCADD severe n=6 median (range) | VLCADD mild n=6 median (range) |
|---|---|---|---|---|---|---|---|
| TG | Saturated lipids⁺ | 15 (10-18) | **18 (17-19)*** | **10 (9.3-12)*** | 14 (12-15) | 13 (9.9-17) | 12 (9.0-20) |
| | Monounsaturated lipids⁺⁺ | 61 (52-63) | **67 (65-67)*** | 67 (57-69) | 64 (59-65) | 60 (58-67) | 61 (55-67) |
| | Polyunsaturated lipids⁺⁺⁺ | 25 (19-30) | **15 (14-19)*** | 26 (20-34) | 21 (1-29) | 27 (18-33) | 25 (16-36) |
| TG(O) | Saturated lipids⁺ | 11 (8.8-14) | 13 (11-14) | **7.9 (6.9-9.7)*** | 11 (10-12) | 11 (7.0-13) | 12 (8.4-12) |
| | Monounsaturated lipids⁺⁺ | 52 (44-56) | **62 (58-63)*** | 50 (43-59) | 50 (44-53) | 48 (45-59) | 49 (45-56) |
| | Polyunsaturated lipids⁺⁺⁺ | 37 (32-46) | **28 (25-29)*** | 43 (33-45) | 38 (36-45) | 42 (30-48) | 39 (32-46) |
| CE | Saturated lipids⁺ | 1.3 (1.0-1.6) | 0.8 (0.7-1.6) | **0.7 (0.6-1.1)*** | 1.3 (1.3-1.9) | 1.0 (0.6-1.4) | 1.2 (0.4-1.4) |
| | Monounsaturated lipids⁺⁺ | 19 (13-20) | 11 (9.7-24) | 16 (13-31) | 24 (22-26) | 16 (12-25) | 21 (8.9-24) |
| | Polyunsaturated lipids⁺⁺⁺ | 79 (70-86) | 88 (75-90) | 83 (68-87) | 75 (72-77) | 83 (74-88) | 78 (75-91) |

| Lipids | Saturation (% of sum) | Controls n=6 median (range) | CPT2D n=3 median (range) | LCHADD n=5 median (range) | MPTD n=3 median (range) | VLCADD severe n=6 median (range) | VLCADD mild n=6 median (range) |
|---|---|---|---|---|---|---|---|
| TG | Short chain lipidsˣ | 63 (55-69) | **74 (69-73)*** | 60 (55-68) | 67 (60-67) | 63 (56-70) | 64 (56-73) |
| | Long chain lipidsˣˣ | 22 (17-27) | 16 (15-20) | 24 (17-29) | 19 (19-25) | 22 (17-29) | 21 (15-28) |
| TG(O) | Short chain lipidsˣ | 53 (47-57) | **61 (59-63)*** | 49 (44-56) | 54 (48-56) | 51 (47-60) | 54 (43-58) |
| | Long chain lipidsˣˣ | 34 (30-40) | **28 (27-30)*** | 39 (31-44) | 34 (32-39) | 36 (28-42) | 33 (30-44) |
| CE | Short chain lipidsˣ | 1.8 (1.7-1.9) | 1.1 (1.0-2.4) | **1.3 (0.8-1.6)*** | 1.7 (1.7-2.6) | 1.5 (0.9-2.0) | 1.6 (0.5-1.9) |
| | Long chain lipidsˣˣ | 76 (70-85) | 86 (70-86) | 82 (73-85) | 73 (66-73) | 80 (70-87) | 75 (70-89) |

**B. PC/PE ratios**

| | Controls n=6 median (range) | CPT2D n=3 median (range) | LCHADD n=5 median (range) | MPTD n=3 median (range) | VLCADD severe n=6 median (range) | VLCADD mild n=6 median (range) |
|---|---|---|---|---|---|---|
| PC/PE ratio | 4.7 (3.6-6.2) | 4.8 (4.6-5.5) | 4.4 (4.3-5.0) | 4.7 (4.3-4.8) | 4.8 (4.0-5.1) | 4.4 (3.6-5.1) |
| D9-PC/D9-PE ratio | 5.4 (4.6-5.7) | 5.6 (4.9-5.8) | 4.9 (4.3-5.4) | 4.8 (4.7-5.6) | 5.1 (4.6-5.4) | 4.4 (4.2-5.8) |

Abbreviations: TG: triacylglycerols, TG(O): alkyldiacylglycerols, CE: cholesterol esters, PC: phosphatidylcholine, PE: phosphatidylethanolamine.
*** [in bold type]** $p < 0.05$, Mann-Whitney U Test compared to controls, ⁺ No double bonds, ⁺⁺ 1-3 double bonds for TG and TG(O) and 1 double bond for CE, ⁺⁺⁺ > 3 double bonds for TG and TG(O) and > 1 double bond for CE, ˣ < 54 carbons for TG and TG(O) and < 18 carbons for CE, ˣˣ >54 carbons for TG and TG(O), and > 18 carbons for CE.

**Supplementary Table 2. Patient plasma samples for confirmation of LPC(14:1)**

| VLCADD ID | lcFAO flux* | Enzyme activity** VLCAD |
|---|---|---|
| LPC_1_ | NA | 19.6% |
| LPC_2_ | NA | 38.5% |
| LPC_3_ | 10 | 3% |
| LPC_5_ | NA | 17.8% |
| LPC_6_ | 33 | 12.3% |
| LPC_7 _MVLCADD_3*** | 39.4 | 7.1% |
| LPC_8MVLCADD_1*** | 50.5 | 9.5% |
| LPC_9 | 87 | 24.1% |
| LPC_10 | 29 | 3.6% |

*lcFAO flux was measured in fibroblasts and is calculated as % of the mean activity in the control fibroblasts analysed in the same experiment. The cell lines were analysed in two independent experiments and the measurements within the experiment were performed at least in duplicate.
**Enzyme activity was measured in fibroblasts and the mean of technical duplicates was expressed as % of the mean of the reference range (1.84-4.80 nmol/(min•mg protein)). NA; not available.
***Patient samples also reported in Table 1, MVLCADD_3 and MVLCADD_1 respectively.

**Supplementary Table 1. Distribution of saturation/ and chain length of neutral lipids and PC/PE ratios**

A. The proportion of neutral lipids (TG, TG(O) and CE) with saturated, mono-unsaturated, and the proportion of short/long neutral lipids are shown as a percentage of the total abundance of the lipid species. Neutral lipids were considered 'short chain' when the average chain length was below 18 (thus: <18 for CE (1 chain), <36 for DG (2 chains), <54 for TG (3 chains), and long-chain with an average chain length of 18 and more. B. The PC/PE ratios in fibroblasts cultured in normal medium and after adding D9-C18:1 to the medium.

**Supplementary Table 2. Enzyme characteristics of patients included for confirmation of LPC(14:1) as VLCADD biomarker**

LPC(14:1) was measured in blood plasma of 8 VLCADD patients with varying disease severity and 11 healthy controls including 1 carrier of a variant in *ACADVL*. Genotype was not available for all cases, lcFAO flux and enzyme activity was included when it has been performed in the included samples.

**Supplementary Figure 1. Correlations of lcFAO-flux with D9-C6:0-carnitine / D9-C18:2-carnitine ratio and D9-20:2-carnitine accumulation**

A. The correlation between the lcFAO flux, a validated diagnostic tool to measure the flux of oleic acid through the β-oxidation system and a good predictor of the VLCADD phenotype and the D9-C6:0-carnitine / D9-9-enoyl-C18-carnitine ratio, representing the β-oxidation capacity. B. The negative correlation between the lcFAO flux and the levels of D9-20:2-carnitine accumulation in LCHADD and MTPD.

**Supplementary Figure 2. Sum of double unsaturated acylcarnitines and very long-chain acylcarnitines in the different LcFAOD.**

A. The sum of the relative abundances of double unsaturated acylcarnitines (C18:2 to C22:2) in lcFAOD patient fibroblasts cultured in standard medium. B. The sum of the relative abundances of very long-chain acylcarnitines (C22:0 to C26:0-carnitine) in lcFAOD patient fibroblasts cultured in normal medium. *= p-value <0.05. **=p-value <0.01. Median (bars) and individual values (black circles) are shown in the graphs.

**Supplementary Figure 3. Eluting peak of OH-C18:1-carnitines**

Extracted-ion chromatogram and bar graphs of OH-C18:1-carnitine and D9-OH-C18:1-carnitine in lcFAOD patient fibroblasts.

**Supplementary Figure 4. Relative abundance of diacylglycerols and alkylacylglycerols.** A. Relative abundance of diacylglycerols in controls and lcFAOD patient fibroblasts cultured in standard medium and D9-C18:1-supplemented medium. B. Relative abundance of alkylacylglycerols in controls and lcFAOD-patient fibroblasts cultured in standard medium and D9-C18:1-supplemented medium. *= p-value <0.05. **=p-value <0.01. Median (bars) and individual values (black circles) are shown in the graphs.

**Supplementary Figure 5. Phospholipids and lysophospholipids**

A. The relative abundance of (lyso)phosphatidylcholine and (lyso)phosphatidylethanolamine in lcFAOD patient fibroblasts and control fibroblasts under standard culture conditions and after adding D9-C18:1 to the medium. B. Relative abundance of LPC in lcFAOD patient fibroblasts and control fibroblasts under standard culture conditions and after adding D9-C18:1 to the medium. *= p-value <0.05. **=p-value <0.01. Median (black lines) and individual values (black circles) are shown in the graphs.

**Supplementary Figure 6. Fragmentation analysis of lipid cluster**

Fragmentation analysis of the lipid cluster in the positive ion mode A. Extracted-ion chromatogram of three high resolution m/z within the unit mass resolution collection window of the quadrupole (left panel). The right panel shows the mass spectrum from m/z 332.0 to m/z 332.5 corresponding to the retention time (RT) indicated by the orange bar in the extracted-ion chromatograms in the left panel. The unknown compound of interest thus eluted at RT 6.81 min and had m/z 332.2618, whereas the other peaks had lower or higher retention times but partly eluted within the window used for the fragmentation analysis. B. MS/MS spectrum of m/z 332.2615 collected using higher energy collisional dissociation (HCD) with a normalized collision energy of 10, 35 and 60 eV where peaks indicated with a red dot elute at RT 6.81 as demonstrated in C. which show the extracted-ion chromatograms of m/z fragments highlighted with a red dot in B. confirming these are fragments of the unknown compound of interest D. Proposed structure of the unknown compound of interest: S-(3-hydroxypalmitoyl)cysteamine, where all red dot fragments from B/C match the proposed structure. The most relevant fragments and their likely chemical formulas are indicated in the table and in the structure. The other fragments mostly represent fragmentation of the aliphatic side-chain.

**Supplementary Figure 7. Excluding extracellular formation of hydrxoyacylcysteamines by methanol/chloroform fixation before sonication**

After adding variable concentrations of D9-C18:1 to the medium (0 to 50 μM), (D9-)C18:1-hydroxycysteamines (A), (D9-)C16:1-hydroxycysteamines (B) and D9-C18:1-hydroxycarnitine were detected in increasing amounts detected in LCHADD patient derived fibroblasts, but not in control fibroblasts. After treatment with methanol:chloroform before sonication, S-(3-hydroxyacyl)cysteamines and hydroxyacylcarnitines were still detected as increased in LCHADD patient derived fibroblasts compared to control fibroblasts, excluding extracellular formation as a consequence of sample preparation. Addition of 25 μM L-carnitine increased formation of both (D9-)S-3(hydroxyacyl)cysteamins (A-B) and D9-hydroxyacylcarnitines (C). Abbreviations: MeOH, methanol
